# Engineered Dendritic Cell-Directed Concurrent Activation of Multiple T cell Inhibitory Pathways Induces Robust Immune Tolerance

**DOI:** 10.1101/706937

**Authors:** Radhika R. Gudi, Subha Karumuthil-Melethil, Nicolas Perez, Gongbo Li, Chenthamarakshan Vasu

## Abstract

Inhibitory/repressor-receptors are upregulated significantly on activated T cells, and have been the molecules of attention as targets for inducing immune tolerance. Induction of effective antigen specific tolerance depends on concurrent engagement of the TCR and one or more of these inhibitory receptors. Here, we show, for the first time that dendritic cells (DCs) can be efficiently engineered to express multiple T cell inhibitory ligands, and enhanced engagement of T cell inhibitory receptors, upon antigen presentation, by these DCs can induce effective CD4+ T cell tolerance and suppress autoimmunity. Compared to control DCs, antigen presentation by DCs that ectopically express CTLA4, PD1 and BTLA selective ligands (B7.1wa, PD-L1, and HVEM-CRD1 respectively) individually (mono-ligand DCs) or in combination (multi-ligand DCs) causes an inhibition of CD4+ T cell proliferation and pro-inflammatory cytokine response, as well as increase in Foxp3+ Treg frequency and immune regulatory cytokine production. Administration of self-antigen (mouse thyroglobulin; mTg) loaded multi-ligand DCs caused hyporesponsiveness to mTg challenge, suppression of autoantibody production, and amelioration of experimental autoimmune thyroiditis. Overall, this study shows that engineered DC-directed enhanced concurrent activation of multiple T cell coinhibitory pathways is an effective way to induce self-antigen specific T cell tolerance to suppress ongoing autoimmunity.

## Introduction

Cytotoxic T-lymphocyte-associated antigen 4 (CTLA4), Programmed cell death-1 (PD1) and B- and T-lymphocyte attenuator (BTLA) are the major inhibitory/repressor/negative-regulatory receptors expressed on T cells, upon T-cell activation in particular^1^. CTLA4 can not only down-regulate T cell responses upon binding to B7.1 and B7.2 (shared ligands of CD28), its interaction with these ligands also curbs activating signals from APCs^2,3^. This receptor, through multiple mechanisms, plays a critical role in peripheral T cell tolerance, T cell homeostasis and regulatory T cell (Treg) function^4–12^. CTLA4 deficient mice undergo a massive lymphoproliferative condition and multi-organ autoimmunity^13,14^ suggesting that this receptor functions as a master regulator of peripheral tolerance mechanism(s). PD1 is a member of the CD28/CTLA4 family, and it is induced on peripheral T and B cells upon activation and its interaction with B7H1 (PD-L1) and B7DC (PD-L2) activates negative regulatory function^15,16^. The ability of PD-L1 on T cells to act as a negative regulator, upon its interaction with B7.1 of APCs, has also been described^17^. PD1 deficiency leads to increased susceptibility to autoimmunity as well as spontaneous autoimmune features at older ages^18^, indicating its contribution to immune tolerance. Very low level of CTLA-4 and PD1 are sufficient for inhibition of the earliest stages of T cell activation through ligand-binding, and co-ligation of these receptors with TCR/CD28 is necessary for this inhibitory function^15,19,20^. Importantly, studies have shown that signaling through CTLA4 and PD1 contributes to the generation and/or functioning of natural and adaptive/induced Tregs^10,21,22^. BTLA, another CD28/CTLA-4 family member, is expressed on activated T and B cells, and its signals can also suppress T-cell response^23^. BTLA interacts with a TNFR family member, HVEM, a herpes virus entry mediator^24,25^. Although HVEM is also known to interact with the T cell activation receptor LIGHT, it dominantly engages BTLA and CD160 on activated T cells through its CRD1 region, and negatively regulates T cell response^25,26^. Although BTLA deficient mice fail to show spontaneous autoimmune features, they do exhibit an increased susceptibility to autoimmunity^23,27^.

Above described aspects suggest that CTLA4, PD1 and BTLA can, individually or in combinations, be excellent targets for promoting peripheral immune tolerance, and the T cell negative regulatory properties of these receptors can be harnessed for autoimmune therapy. Importantly, inhibition of T cell response requires delivering active negative signals through these repressor receptors in conjunction with TCR engagement^19,28–31^. Therefore, induction of antigen specific tolerance by targeting CTLA4, PD1 and/or BTLA may require concomitant engagement of antigen specific TCRs and enhanced signaling through these receptors. Previously, we have shown that dominant engagement of CTLA-4 on T cells from target tissues and antigen presenting dendritic cells (DCs) can induce Treg response, antigen specific T cell tolerance, and prevent and treat autoimmune experimental autoimmune thyroiditis (EAT) and spontaneous type 1 diabetes (T1D) in mouse models^32–36^. We have also shown that dominant engagement of CTLA-4 by its preferential ligand, B7.1 on DCs could lead to the induction of adaptive Tregs and suppression of T1D^34^. Previous reports have also shown that DCs that are depleted of co-stimulatory molecules induce tolerance^37,38^, Nevertheless, exogenous expression of ligands in primary immune cells such as DCs for selectively enhancing T cell inhibitory receptor engagement, concurrent targeting of multiple receptors particularly, for generating tolerogenic antigen presenting cells (tAPCs) has not been reported before. Here, we tested if “potent” tAPCs can be generated by engineering the DCs to exogenously express selective ligands for CTLA4, PD1 and BTLA individually (mono-ligand DCs) or in combination (multi-ligand DCs) to harness the T cell inhibitory functions of these receptors and produce robust immune tolerance. We show that DCs can be efficiently engineered to simultaneously express multiple T cell repressor receptor-selective ligands using a lentiviral transduction approach, and antigen-loaded ligand-expressing engineered DCs, multi-ligand DCs in particular, can act as powerful tAPCs. These engineered DCs can induce profound inhibition of T cell proliferation, modulation of cytokine response, and generation of T cells with a regulatory phenotype, both in vitro and in vivo. Treatment of mouse model of experimental autoimmune thyroiditis (EAT) using mouse thyroglobulin (mTg)-loaded inhibitory ligand-DCs caused suppression of autoreactive T cell and antibody responses and amelioration of thyroiditis severity. Overall, this study, for the first time, demonstrates that engineering the DCs for expressing ligands specific for multiple T cell inhibitory receptors concomitantly is an effective way to produce tAPCs, and enhanced activation of multiple T cell repressor pathways by these cells can effectively suppress the ongoing autoimmunity.

## Results

### T cell negative regulatory ligand expressing DCs modulate T cell response in vitro

Since negative regulatory pathways in T cells play a key role in peripheral immune tolerance, we hypothesized that overexpression of selective ligands for CTLA4, PD1 and BTLA receptors could make DCs produce tolerogenic effects on T cells through enhanced receptor engagement upon antigen presentation. To test this hypothesis, we engineered the bone marrow (BM) derived DCs to express the ligands for them, specifically B7.1wa, PD-L1 and HVEM-CRD1 exogenously by employing a lentiviral transduction system (Supplemental Figs. 1 and Fig. 1A) and characterized the phenotypic and functional properties of these cells. First, we determined if lentiviral transduction alone affects the viability and functional properties of DCs. While both untreated and LPS treated lentiviral transduced DCs retained phagocytic ability (Fig. 1B) and viability (Supplemental Fig. 2), virus transduced, untreated cells expressed relatively higher levels of costimulatory molecules (Fig. 1C) and cytokines (Fig. 1D), and induced greater proliferation and cytokine production in T cells upon antigen presentation (Figs. 1E & 1F) suggesting that lentiviral transduction alone could activate the DCs. On the other hand, LPS treated non-transduced and lentivirus transduced DCs produced more or less comparable levels of surface markers and cytokines. Further, compared to non-treated DCs, LPS-treated non-transduced and virus-transduced DCs induced comparable, although significantly higher than untreated DCs, degrees of T cell activation and cytokine production. Therefore, since effective initial activation of T cells is an important step in inducing Ag specificity to T cell tolerance and to minimize susceptibility to changes in their antigen presentation properties in vivo when injected, we used LPS exposed control and ligand expressing DCs for rest of the study.

**FIGURE 1:**
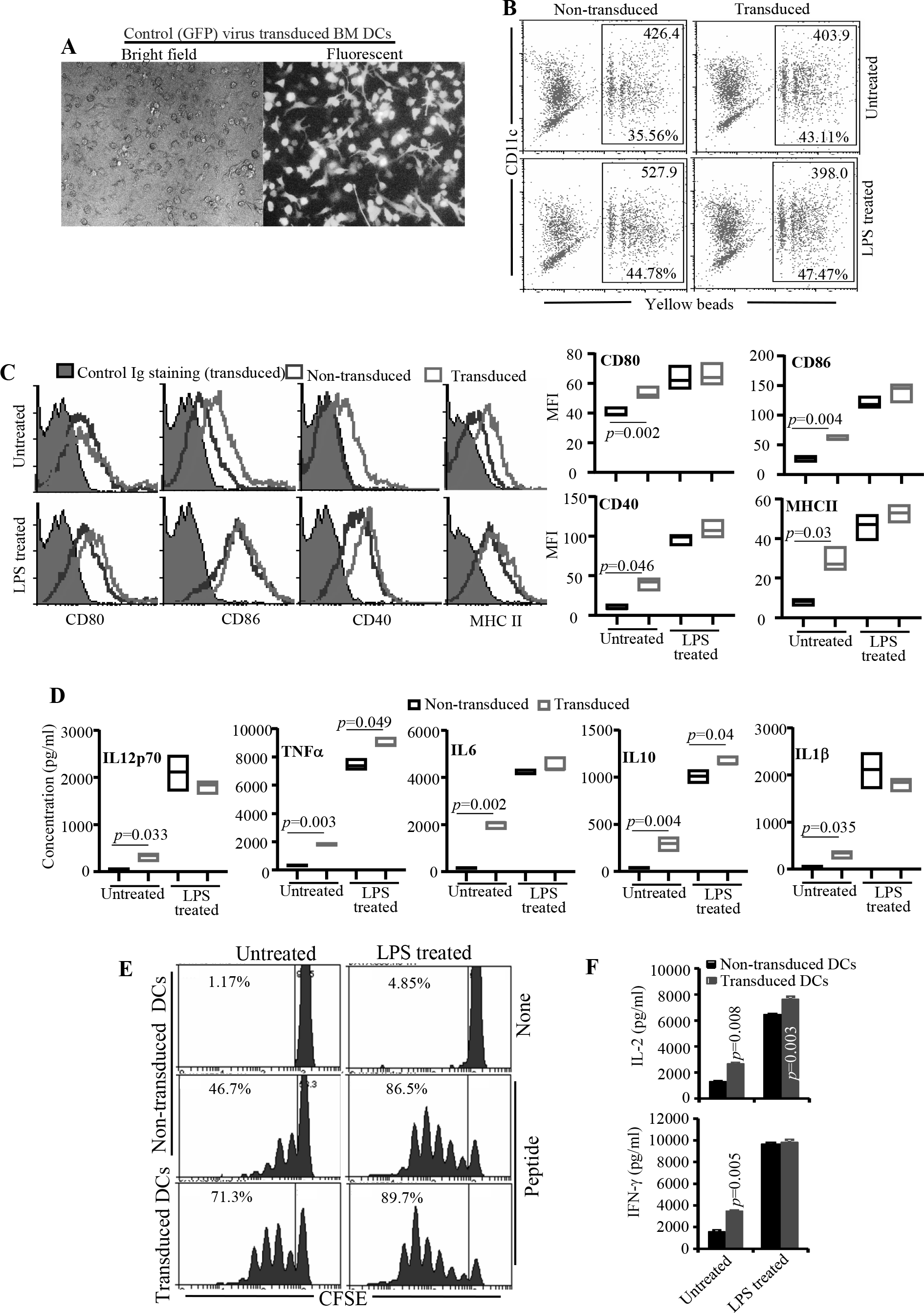
Characterization of antigen presentation related properties of lentivirus transduced DCs. C57/BL6 BM DCs were transduced with lentiviral vectors as described in Materials and methods. To induce maturation, lentivirus transduced and non-transduced cells were incubated with bacterial LPS (1 μg/ml) for 24 h. **A)** An example of transduction of BM DCs using control (GFP) virus. **B)** Lentivirus transduced and non-transduced DCs were subjected to phagocytosis assay. Cells were incubated with 1 μm yellow fluorescent beads for 2 h, stained for CD11c and analyzed by FACS. Percentage of cells positive for yellow fluorescence and mean (yellow) fluorescence value (MFI) of gated population from a representative experiment are shown. For fluorescence compatibility, CFP vector transduced DCs were used for this assay. **C)** Lentivirus transduced and non-transduced cells were examined for the expression levels of antigen presentation related activation markers by FACS. Representative FACS plots (left panel) and Mean±SD of MFI values of cells from 3 independent, parallel, transductions (right panel) are shown. Both virus transduced and non-transduced cells were also stained using control Ig, but showed only the histograms of transduced DCs as representative background staining. **D)** Supernatants of virus transduced and non-transduced cells (2× 10^6^ cells/ml) described for panel C were collected from 24h culture and examined for cytokine levels by Luminex multiplex assay. Mean values of samples from 3 independent, parallel, transductions, each tested in triplicate, are shown. *p*-value was calculated by paired *t*-test. **E)** Ova peptide pulsed virus transduced and non-transduced DCs were used in antigen presentation assays by culturing with CFSE labeled purified CD4+ T cells from OT-II mice. CFSE dilution was determined by FACS on day 4 after staining with PE-labeled anti-CD4 Ab. CD4+ cells were gated for the shown histograms. Representative FACS plots of 3 independent, parallel, experiments done in triplicate are shown. **F)** Supernatants obtained from the above cultures on day 4 were tested for IL2 and IFNγ levels by ELISA. Mean±SD values of 3 independent experiments, done in triplicate, using parallel preparations of DCs are shown. *p*-value by paired *t*-test. Lentivirus transduced DC group was compared to respective non-transduced DC group. This assay was repeated at least twice with similar trends.

To assess the functional ligand expression, B7.1wa, PD-L1 and HVEM-CRD1 lentivirus transduced DCs (Fig. 2A) were examined for ligand levels and the soluble receptor binding abilities by FACS. Staining using ligand specific antibodies showed that virus transduced cells express profoundly higher levels of respective ligands compared to non-transduced cells (Fig. 2B). Further, incubation with soluble receptors followed by fluorochrome-labeled secondary antibody reagent demonstrated greater degree of binding by respective receptors in transduced cells compared to control cells, indicating that these exogenously expressed ligands are functional in terms of engaging the respective receptors. Of note, since WT B7.1 binds to both CTLA4 and CD28, and WT HVEM binds to BTLA and LIGHT, previously described CTLA4 and BTLA selective binding abilities of B7.1wa and HVEM-CRD1^39,40^, were confirmed using 293T cells expressing these ligands (not shown).

**FIGURE 2:**
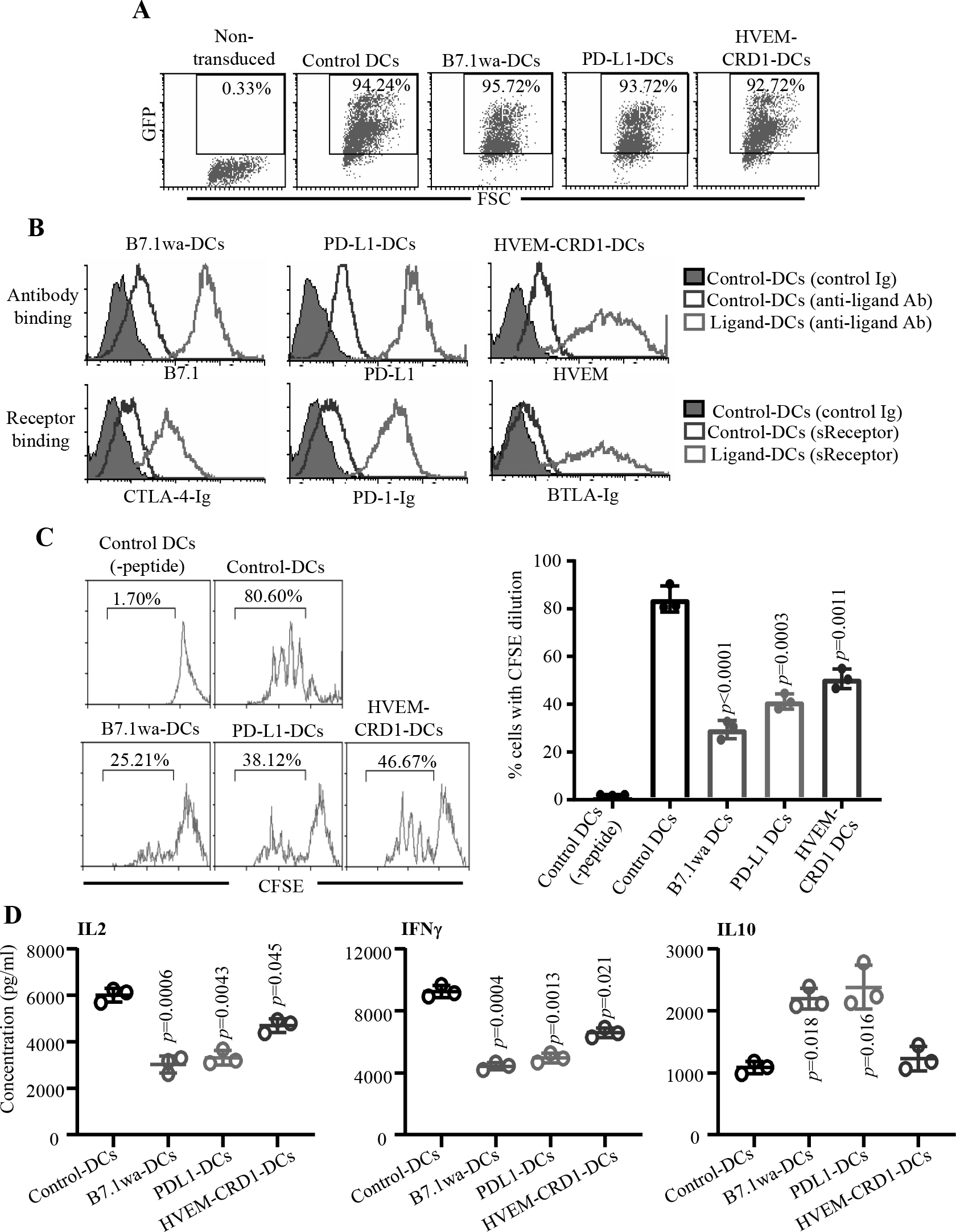
T cell negative regulatory ligand expressing engineered DCs modulate T cell response in vitro. C57/BL6 BM DCs were transduced with control, B7.1wa, PD-L1, and HVEM-CRD1 lentivirus (GFP under an independent promoter) for 48 h and treated with bacterial LPS for 24 h. **A)** Control and ligand engineered cells were examined for GFP expression by FACS. Representative FACS graphs (and percentage GFP +ve cells) from multiple transductions are shown. **B)** Ligand expression levels were determined after staining with specific Abs (upper panels). Cells were also incubated with soluble receptors (CTLA4-Ig, PD1-Ig or BTLA-Ig), followed by PE-labeled Fab(2) fragment of anti-IgG (Fc specific) Ab and tested by FACS (lower panels). Histogram overlay graphs (representative of 3 independent experiments) showing ligand specific staining of control virus transduced and ligand virus transduced DCs are shown. Both control and ligand DCs were also stained using control Ig reagents, but showed only the histograms of control DCs as representative background staining. **C)** OT-II peptide pulsed control and ligand engineered DCs were used in antigen presentation assays by culturing with CFSE labeled purified CD4+ T cells from OT-II mice. CFSE dilution was determined on day 4 by FACS, after staining for CD4. CD4+ cells were gated for the histograms shown here. **D)** Supernatants obtained from these cultures on day 4 were tested for IL-2 by ELISA. Equalized number of cells from the primary cultures were also stimulated using Ova peptide pulsed freshly isolated splenic DCs for 24 h and the supernatants were tested for IFNγ and IL10 levels by ELISA. Mean±SD values of 3 independent experiments, done in triplicate, using parallel preparations of DCs are shown for panels C and D. *p*-value by paired t-test. Each ligand DC group was compared separately with control DC group. This assay was repeated at least twice with similar trends.

To examine differences in the abilities of T cell negative regulatory ligand expressing DCs to activate T cells, control, B7.1wa, PD-L1 and HVEM-CRD1 DCs were pulsed with OT-II peptide and cultured with CD4+ T cells enriched from OT-II mouse spleen. As observed in Fig. 2C, T cells cultured with all three ligand DCs showed an overall lower rate of proliferation as compared to those cultured with control engineered DCs. Importantly, to determine if the suppression of T cell proliferation is due to enhanced receptor engagement, anti-CTLA4 antibody, PD1-Ig and BTLA-Ig were added to respective cultures to block receptor-ligand interaction. Supplemental Fig. 3 shows that blockade of receptor interaction with respective ligand in the culture reversed the suppressive effect on T cell proliferation induced by selective ligand expressing engineered DCs, supporting the notion that ligand DC induced effects are primarily due to enhanced T cell inhibitory receptor engagement.

To further assess the effect of enhanced T cell inhibitory receptor engagement upon antigen presentation, supernatants from primary cultures were examined for IL2 levels. CD4+ T cells activated using all three inhibitory ligand DCs produced significantly lower amounts of IL2 compared to control DC activated cells (Fig. 2D). Importantly, analysis of cytokines secreted by T cells from these primary cultures, upon antigen challenge, shows that antigen presentation by ligand DCs, B7.1wa-DCs and PD-L1-DCs particularly, induced IL10 producing T cells. Further, T cells activated using all three ligand DCs produced significantly lower amounts of IFNγ compared to control DC activated T cells. Overall, these observations suggest that enhancing the CTLA4, PD1 and BTLA selective ligand strength on antigen presenting DCs results in suppression of T cell responses, albeit different outcomes in-terms of the type of response when different receptors were targeted.

### Multi-ligand DCs suppress T cell responses more effectively than mono-ligand DCs in vitro

Next, we examined if multiple T cell negative regulatory ligands can be efficiently expressed simultaneously on DCs and it provides the DCs with a better ability, as compared to expression of individual ligands, to suppress or modulate T cell response upon antigen presentation. To test this, multi-ligand DCs were generated using lentiviral vectors encoding different fluorescent-tag versions of T cell negative regulatory ligands (B7.1wa-CFP, PD-L1-YFP and HVEM-CRD1-RFP; shown in Supplemental Fig. 1B), and the ligand expression levels and receptor binding abilities were confirmed microscopically and/or by FACS (Fig. 3A and 3B). The multi-ligand DCs were then, along with mono-ligand and control DCs, pulsed with OT-II peptide and used for antigen presentation assays in vitro. As observed in Fig. 3C and 3D, fluorescent-tag mono-ligand DCs induced proliferative and IL2 responses in OT-II CD4+ T cells, similar to the response induced by non-tagged ligand DCs (shown in Fig. 2). Importantly, the proliferative and IL2 responses of OT-II CD4+ T cells were significantly lower when multi-ligand DCs, compared to individual ligand or control DCs, were used for antigen presentation.

**FIGURE 3:**
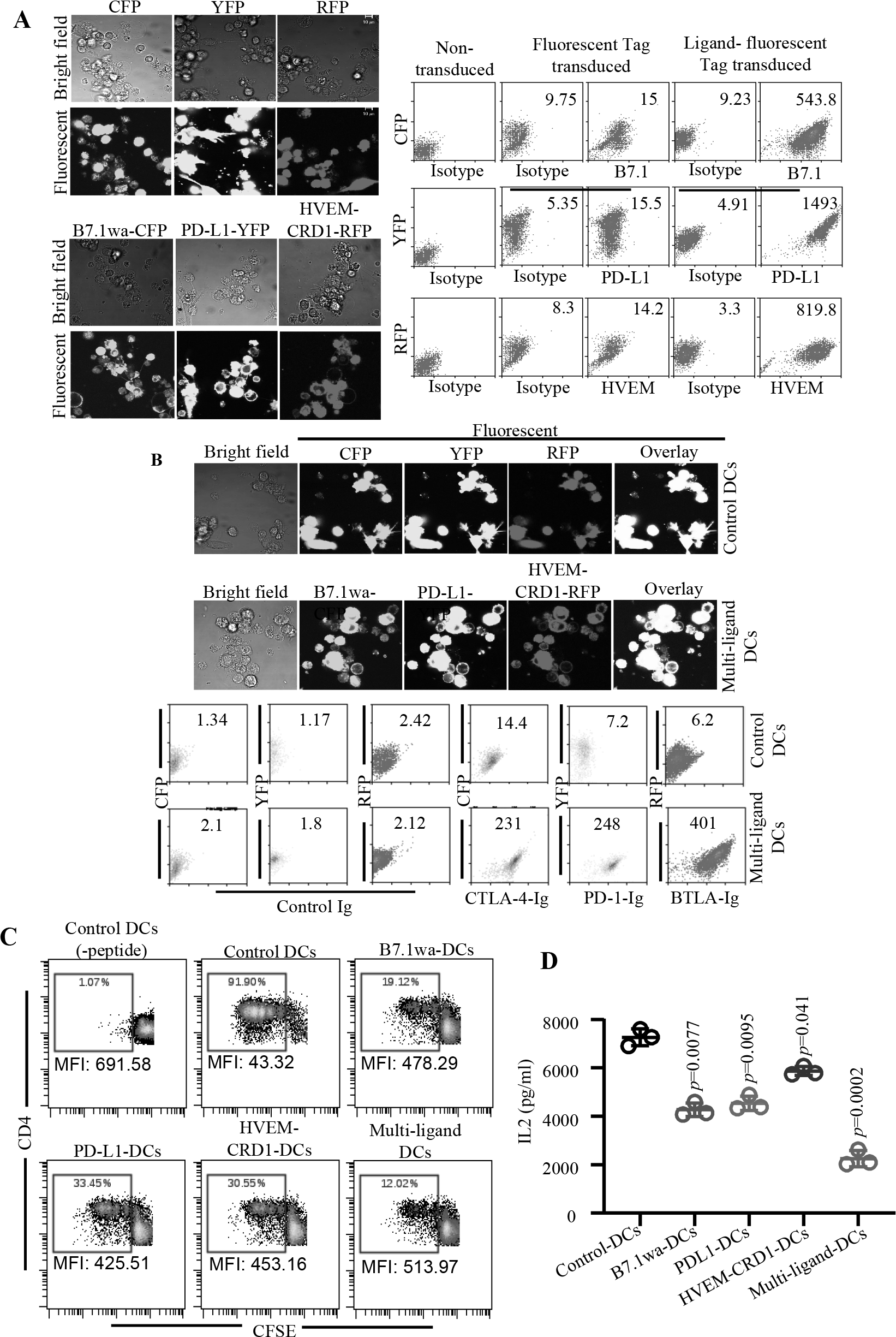
Multi-ligand DCs suppress T cell response more effectively than mono-ligand DCs in vitro. BM DCs were transduced with lentiviral particles generated using CFP, YFP, RFP, B7.1wa-CFP, PD-L1-YFP, and HVEM-CRD1-RFP plasmids for 48 h and treated with bacterial LPS for 24 h as described under Materials and methods. **A)** Cells were transferred to coverslip bottom chamber slides and incubated overnight, and subjected to confocal microscopy to detect fluorescent protein expression (left panel). Cells were also tested for ligand expression after staining with B7.1, PD-L1 and HVEM specific Abs (right panel) by FACS. **B)** Cells were transduced with mixtures of all three control vectors or ligand vectors, and processed for microscopy (upper panels) and FACS after staining using soluble receptors followed by anti-IgG (Fc specific) Ab (lower panels). Mean fluorescence index (MFI; log scale) values for ligand specific staining are shown for both panels A and B. Representative graphs of multiple transductions are shown. **Note:** Lack of correlation between fluorescence tags and ligand specific staining (using either the antibody or soluble receptor) indicates the detection limitation of FACS for these tags, CFP and RFP particularly. **C)** Ova peptide pulsed control and ligand engineered DCs were used in antigen presentation assays by culturing with CFSE labeled purified CD4+ T cells from OT-II mice. CFSE dilution was determined on day 4 by FACS after staining for CD4. CD4+ cells were gated for the graphs shown here. **D)** Supernatants obtained from these cultures on day 4 were tested for IL2 levels by ELISA. Representative FACS plot (for panel C), and mean±SD values of 3 independent experiments, done in triplicate, using parallel preparations of DCs (for panel D) are shown. This assay was repeated at least twice with similar trends. *p*-value by paired *t*-test. Each ligand DC group was compared separately with control DC group.

Of note, while the levels of early activation marker CD69 were relatively lower in individual and multi-ligand DC activated T cells compared to control DC activated cells at 16 h time-point, this difference was considerably increased by 48h time-point (Supplemental Fig. 4A). Similarly, the differences in the degree of suppression of IL2 production by control and ligand DCs was more profound at 72 h, compared to 36h, time-point (Supplemental Fig. 4B). These observations suggest that the early activation of T cells is suppressed less profoundly than later activation state by negative regulatory ligand expressing DCs, and the enhanced inhibitory receptor engagement occurs mainly after initial activation of T cells. It is important to note, as reported before^13,17,19,23,40^, that initial activation of T cells by antigen specific TCR engagement is critical for upregulating the expression of T cell inhibitory receptors.

### Multi-ligand DCs modulate T cell function more effectively than mono-ligand DCs

To further realize the impact of enhanced engagement of repressor receptors by multi-ligand DCs upon antigen presentation, pro- and anti-inflammatory cytokine profiles and Foxp3 expression of CD4+ T cells activated using engineered DCs were determined. Figs. 4A and 4B show that fluorescent-tag individual ligand DCs induce IL10 and IFNγ responses similar to that induced by untagged ligand DCs (shown in Fig. 2). Importantly, multi-ligand DC mediated activation of CD4+ T cells, similar to B7.1wa DC and PD-L1 DC mediated activation, resulted in the generation of profoundly higher frequencies of IL10 producing T cells in the in vitro cultures compared to that by control DCs. Reciprocally, these cultures showed significantly diminished IFNγ producing CD4+ T cell frequencies compared to control DC containing cultures. Interestingly, IL17 response of T cells was higher upon antigen presentation by B7.1wa-DCs, but not by other ligand DCs. Notably, only B7.1wa-DCs and multi-ligand DCs, but not other DCs, induced an increase in Foxp3+ T cell frequencies and active TGF-β1 production upon antigen presentation in these cultures (Figs. 4A and 4B). This suggested that TGF-β1 could be responsible for inducing higher Foxp3+ T cells in an auto/para-crine manner in B7.1wa-DC and multi-ligand DC containing cultures, and higher IL17 production in B7.1wa-DC cultures. This notion has been substantiated by the reduction of Foxp3+ CD4+ T cell frequencies in these cultures (Fig. 4C) and a suppression of IL17 production along with an increase in IFNγ response in B7.1wa-DC cultures (Supplemental Fig. 5) upon addition of TGF-β1 neutralizing antibody. We also determined IL10+ and LAP+ Tregs (Foxp3+) or ‘effector’ (Foxp3-) T cell frequencies in cultures similar to that described for Fig. 4. Although surface LAP expression may not correlate with active TGFβ1 levels in the cultures, as observed in Supplemental Fig. 6, difference in LAP expression was observed primarily with Foxp3(GFP)+ cells of B7.1wa and multi-ligand DC cultures. Further, while IL10 production in B7.1wa and multi-ligand DC cultures appears to be associated with both Foxp3+ and Foxp3-populations, this cytokine in PD-L1 and HVEM-CRD1-DC containing cultures is primarily of Foxp3-CD4+ T cell origin. Overall, these observations show that while all three ligand-DC preparations induce modulation of T cell response, multi-ligand DCs show more profound modulation of T cell function than that induced by individual ligand DCs, and suggest that these cells could be more efficient tAPCs compared to individual ligand DCs.

**FIGURE 4:**
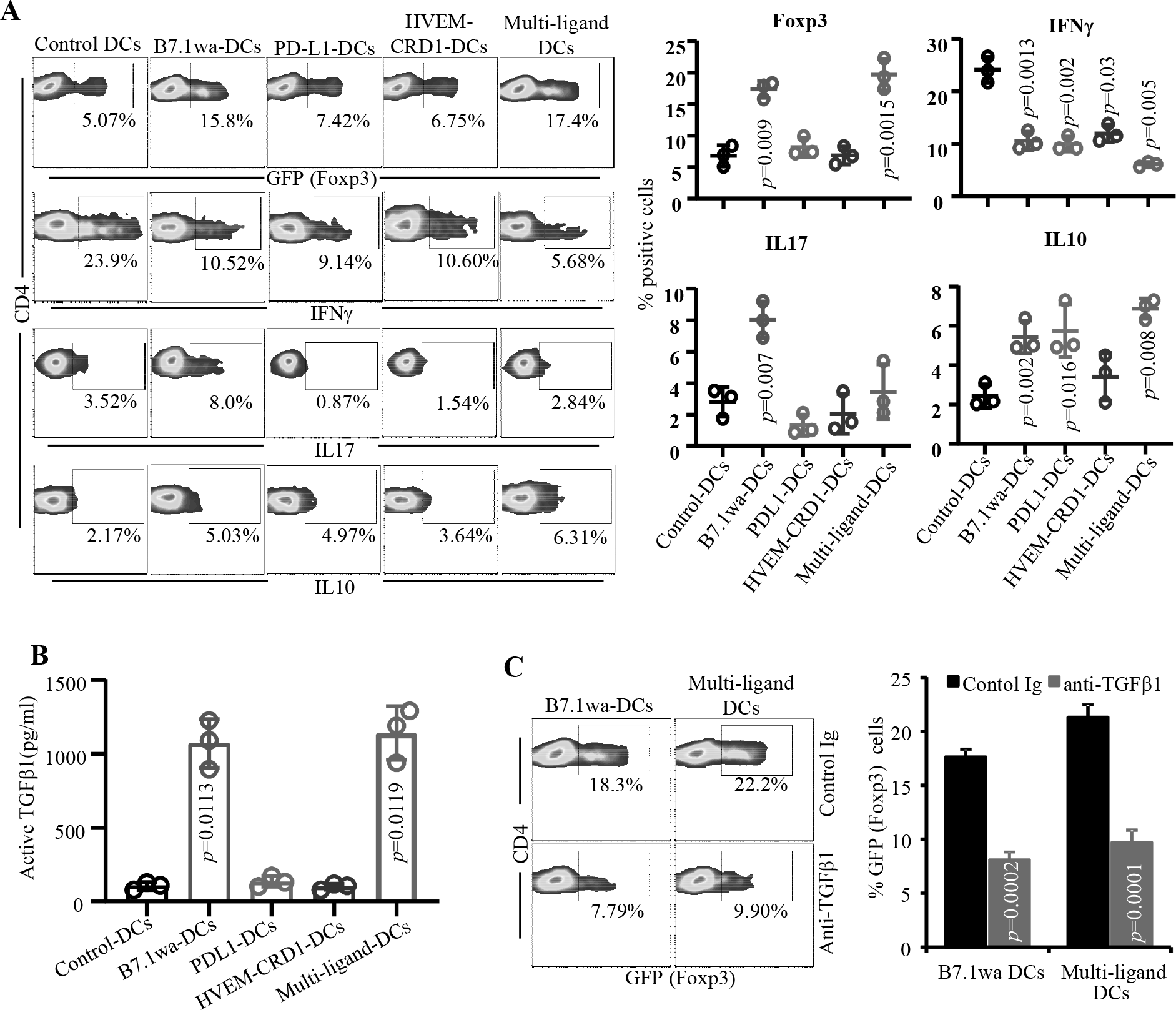
Multi-ligand DCs modulate T cell function more effectively than mono-ligand DCs in vitro. BM DCs were transduced with lentiviral particles generated using CFP, YFP, RFP, B7.1wa-CFP, PD-L1-YFP, and HVEM-CRD1-RFP plasmids, and used in antigen presentation assays as described for Fig. 3. CD4+ cells isolated from OT-II-Foxp3-GFP mouse spleens were used in this assay. **A)** OT-II T cells from similar primary cultures were examined for GFP+ cell frequencies by FACS. Cells from primary cultures were stimulated using PMA and ionomycin in the presence of BrefeldinA for 4 h, stained for intracellular cytokines IFNγ, IL17 and IL10, and subjected to FACS analysis. CD4+ cells were gated for the histograms shown here. Representative FACS plots (left panel) and Mean±SD of % GFP(Foxp3) or cytokine positive cell frequencies of 3 independent experiments, done in triplicate, using parallel preparations of DCs (right panel) are shown. **B)** Supernatants obtained from the above primary cultures were tested for active TGF-β1 levels by ELISA. Mean±SD values of 3 independent experiments, done in triplicate, using parallel preparations of DCs are shown. **C)** Antigen presentation assay using B7.1wa DCs and multi-ligand DCs and OT-II-Foxp3-GFP were conducted in the presence of isotype control antibody or TGF-β1 neutralizing antibody, and examined for GFP(Foxp3)+ T cell frequencies. Representative FACS plot (left panel) and Mean±SD of % GFP (Foxp3)+ cell frequencies of 3 independent experiments, done in triplicate, using parallel preparations of DCs (right panel) are shown. The assay was repeated once with similar trend. *p*-value by paired *t*-test for all panels. Each ligand DC group was compared separately with control DC group.

### Multi-ligand DCs modulate T cell response more effectively than mono-ligand DCs in vivo

To examine the ability of T cell negative regulatory ligand-expressing DCs to modulate antigen specific T cell response in vivo, Ova primed C57BL/6 mice were treated with Ova-pulsed control and ligand DCs and examined for the T cell phenotype and proliferative response, and Ova specific antibody levels. As observed in in vitro assays (shown in Fig. 4), significantly higher frequencies of splenic CD4+ T cells from B7.1wa-DC and multi-ligand DC recipient mice were Foxp3+ compared to control DC recipient mice (Fig. 5A). Furthermore, CFSE dilution assay demonstrated that the abilities of CD4+ T cells from B7.1wa, PD-L1, HVEM-CRD1, and multi-ligand DC recipients to respond to Ova challenge were relatively lower compared to T cells from control DC recipients, with the T cells from multi-ligand DC recipients showing the least proliferation upon challenge with Ova (Fig. 5B). Cytokine response of T cells from ligand DC treated mice showed a trend similar to that observed in the in vitro assay of Fig. 4 (Fig. 5C). Importantly, better suppression of anti-Ova antibody response was achieved when the mice were injected with multi-ligand DCs compared to those that received mono-ligand DCs (Fig. 5D). Overall, these results suggest that while enhanced engagement of different repressor receptors by antigen presenting DCs modulates T cell function differently, they exert more effective regulation of T cell, and eventual B cell, responses upon co-engagement. One of the key common features of all ligand DCs, albeit the differences in magnitudes, appears to be the ability to induce IL10 production in T cells suggesting the contribution of this cytokine to hypo-responsiveness of T cells and to the overall immune tolerance. Hence, we determined the contribution of IL10 to immune modulation mediated by ligand-DC activated CD4+ T cells. As observed in Supplemental Fig. 7A, CD4+ T cells from both WT and IL10−/− mice showed increased frequencies of Foxp3+ cells when activated in the presence of B7.1wa and multi-ligand DCs. Importantly, in vitro T cell co-culture assays showed that IL10 deficiency significantly diminishes the regulatory/suppressive properties of all ligand DC activated T cells (Supplemental Fig. 7B).

**FIGURE 5:**
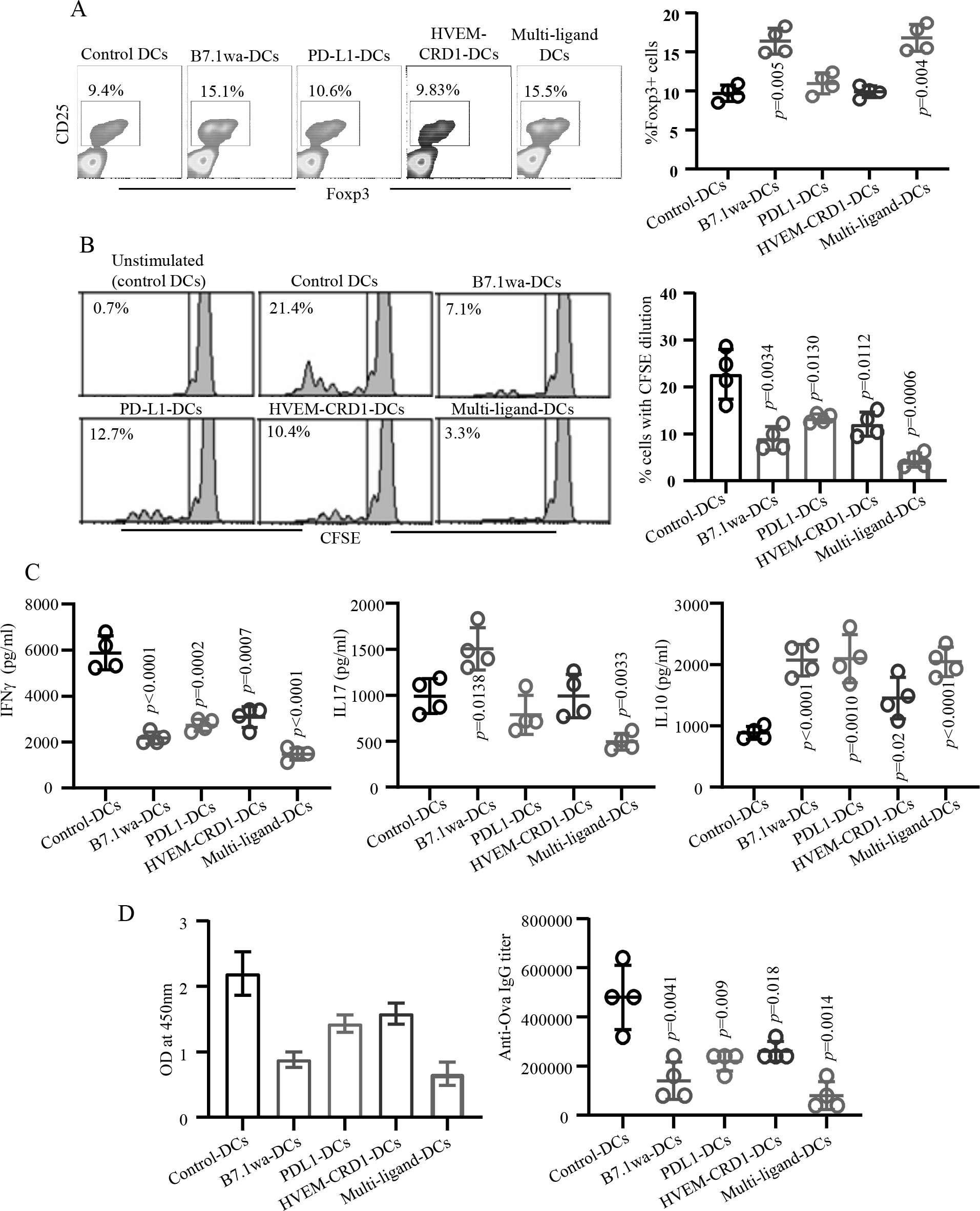
T cell inhibitory ligand expressing engineered DCs modulate T cell response in vivo. OVA primed (days 0 and 10) C57/BL6 mice were treated with OVA pulsed, LPS treated control, B7.1wa, PD-L1, HVEM-CRD1 or multi-ligand DCs (days 5 and 15) as described under Materials and methods. Fluorescent protein tagged constructs were used; control transduction was done using a pool of viruses encoding all three fluorescent proteins as described in Fig. 3. These mice were euthanized at day 25 for examining T cell characteristics. **A)** Spleen cells were examined for Foxp3+ cell frequencies by FACS. **B)** CFSE stained spleen cells were challenged ex vivo with OVA for 4 days, and examined for CFSE dilution by FACS. Representative FACS plots (left panels) and Mean±SD (4 mice/group) of percentage of CD4+ T cells with Foxp3 expression or CFSE dilution (right panels) are shown for panels A and B. Assays were performed using cells from individual mice in triplicate. **C)** Spleen cells were cultured in the presence of Ova for 48 h and the supernatants were tested for cytokine levels by ELISA. Mean±SD values of cytokine levels in 4 mice/group tested in triplicate are shown. **D)** Serum samples from these mice were tested for anti-mTg total IgG antibody levels by ELISA. Mean±SD of OD values (left panel) and antibody titer (right panel) of 4 mice/group are shown. This experiment was repeated once with same number of mice. *p*-value by unpaired *t*-test for all panels. Each ligand DC group was compared separately with control DC group.

### Treatment with T cell negative regulatory ligand expressing DCs during EAT induction results in suppression of autoimmunity and thyroiditis severity

To determine the ability of ligand DCs to modulate autoimmune response in vivo, mTg immunization model of EAT was employed. CBA/J mice were immunized with mTg on days 0 and 10, and cohorts of mice were treated during the disease induction stage with mTg pulsed control or multi-ligand DCs on days 5 and 15, and the degree of thyroiditis severity and the autoantibody response were determined on day 40. As shown in Fig. 6A, while treatment using mono-ligand DCs, compared to control DCs, resulted in significant suppression of thyroiditis severity, mice that received multi-ligand DCs showed profoundly lower thyroiditis severity. Further, while treatment of EAT mice with mono-ligand DCs and multi-ligand DCs resulted in significantly lower mTg specific IgG1 and IgG2a (Fig. 6B) autoantibody production compared to control DC treated mice, with the multi-ligand DC treated group showing the most profound suppression of autoantibody production. Draining cervical LN cells from cohorts of mice euthanized on day 30 were examined for proliferative and cytokine responses upon mTg challenge ex vivo. Ligand DC treated mice, compared to control DC recipients, showed significantly lower T cell proliferative response upon challenge with mTg (Supplemental Fig. 8), a trend similar to that of Ova primed mice described under Fig. 5. Importantly, as observed in Ova primed mice as well as in the in vitro assays, both B7.1wa and multi-ligand DC treated mTg primed mice showed significantly higher frequencies of Foxp3+ CD4+ T cells compared to other groups (Fig. 6C). Interestingly, LN cells from all ligand DC recipient mice produced profoundly lower IFNγ and higher IL10 upon ex vivo challenge with mTg (Fig. 6D). On the other hand, while CD4+ T cells from B7.1wa DC treated mice produced higher amounts of IL17 compared to control DC recipients, IL17 response of T cells was lower when the mice were treated with PD-L1 DCs or multi-ligand DCs. These observations have been substantiated by detection of correlating frequencies of IFNγ+, IL17+ and IL10+ CD4 T cells upon intracellular staining of fresh splenic CD4+ from ligand DC treated mice (Supplemental fig 9). Overall, these results show that T cell negative regulatory ligand expressing DCs, multi-ligand DCs particularly, have the potential to induce robust hyporesponsiveness against self-antigens and promote protection against autoimmune destruction of target organ.

**FIGURE 6:**
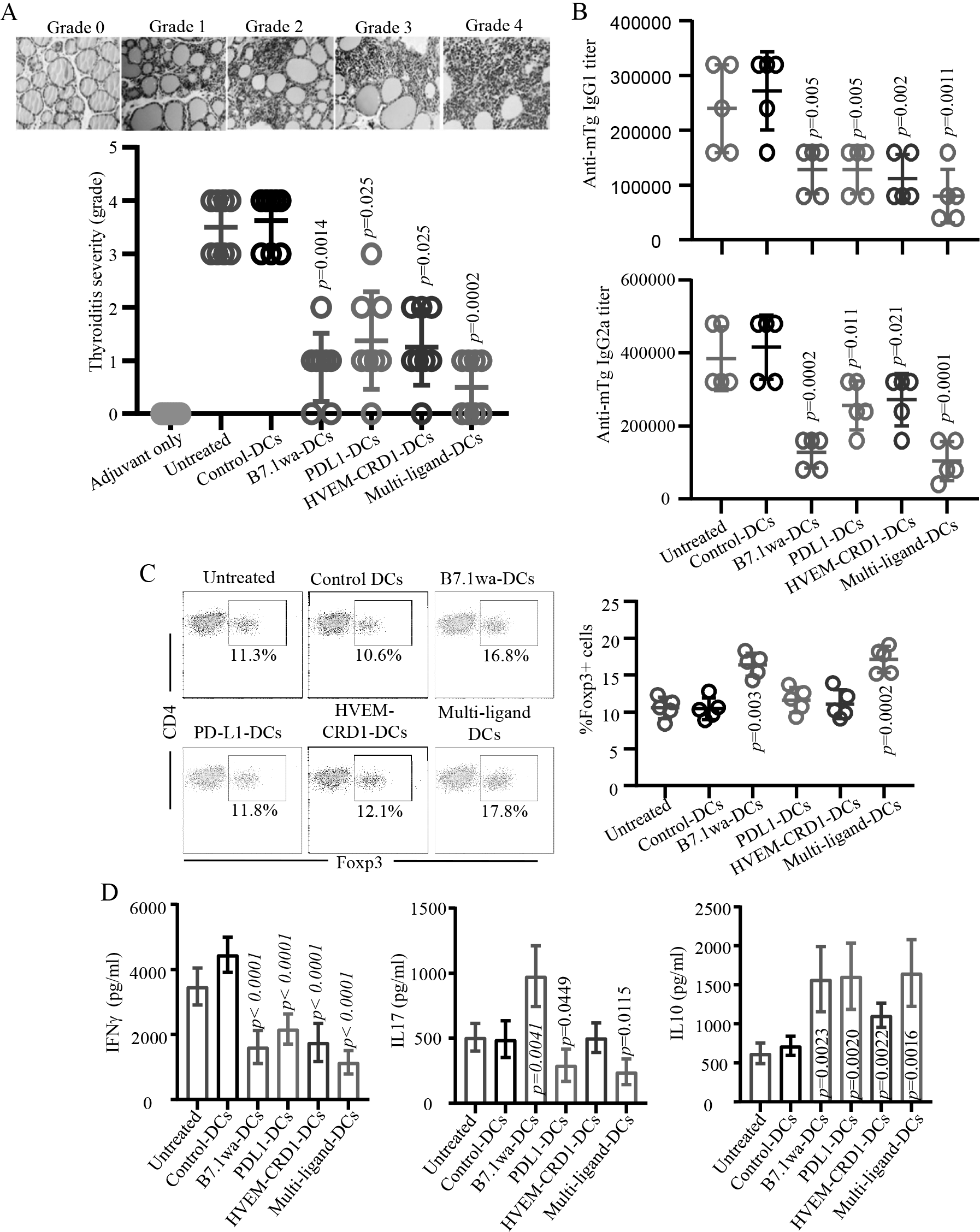
Treatment with T cell negative regulatory ligand expressing DCs results in suppression of EAT. CBA/J mice were immunized i.v. with LPS+PBS (adjuvant control) or with LPS+mTg (on days 0 and 10). Sets of LPS+mTg injected mice were left untreated (untreated control) or treated with mTg pulsed control DCs, mono-ligand or multi-ligand DCs (2×106 cells/mouse/ i.v.) on days 5 and 15. As done for Fig. 5, fluorescent protein tagged constructs were used. Cohorts of mice were euthanized on day 40, thyroids were harvested for determining thyroiditis severity. **A)** H&E stained thyroid sections were examined for immune cell infiltration and follicular destruction. The severity of infiltration and follicular damage was graded, and representative images of different grades of thyroiditis (upper panel) and the percentages of mice (n=8 mice/group) with different thyroiditis severity (lower panel) are shown. *p*-value by Fisher’s exact test comparing mice with thyroiditis grades ≤1 and ≥2 in ligand DC recipients against control DC recipients. **B)** Serum samples were tested for anti-mTg IgG1 and IgG2a antibody levels by ELISA. Mean±SD values of antibody titer of 5 mice/group tested in triplicate are shown. **C)** Cervical LN cells from one cohort of mice from a similar experiment euthanized on day 30 post-mTg priming were examined for Foxp3+ cell frequencies by FACS. Representative FACS graphs (left panel) and mean±SD of Foxp3+ CD4 T cell frequencies of 5 mice/group (right panel) are shown. **D)** Cervical LN cells were cultured in the presence of mTg for 48 h and the supernatants were tested for cytokine levels by ELISA. Mean±SD values of cytokine levels in 5 mice/group tested in triplicate are shown. *p*-value by unpaired *t*-test for panels B to D. Each ligand DC group was compared separately with control DC group.

## Discussion

DCs are considered the most effective APCs and the only type of cells that are capable of activating naïve T cells^41,42^. Hence, they can play an important role in breaking tolerance and in initiating autoimmune responses. These APCs are capable of capturing antigens from the site of inflammation and presenting it to T cells at lymphoid organs^43^. These properties make the DCs most efficient APCs, and indicate that they can be used to effectively induce T cell activation or tolerance if the immunological synapse is appropriately manipulated. While DCs maintained in an immature state are considered tolerogenic due to significantly low expression of costimulatory molecules and/or pro-inflammatory cytokines^44,45^, these cells are highly susceptible to maturation in vivo, especially under pro-inflammatory microenvironment in autoimmune conditions, and can produce adverse effects during therapy. Hence, engineering the DCs to ensure the maintenance of their tolerogenic function, even under the pro-inflammatory conditions, is critical for autoimmune therapy. In this regard, DCs that are selectively deleted of costimulatory molecules or ectopically expressing cytokine and non-cytokine factors have shown the ability to induce T cell tolerance^37,46,47^. On the other hand, we reported that enhancing CTLA4 agonist strength on mature DC surface could lead to dominant engagement of repressor-receptor on T cells upon antigen presentation and induce antigen specific Tregs and T cell tolerance^35,36^. This suggested to us that enhancing the strength of one or more T cell repressor receptor selective ligands by exogenous expression can be an effective strategy to generate efficient tolerogenic DCs and this approach will have immense potential in suppressing autoimmunity. However, exogenous or over-expression of multiple factors in DCs have been challenging due to the difficulty in transfecting these primary immune cells efficiently. This study, for the first time, demonstrates that lentiviral transduction system can be employed to generate tAPCs that are exogenously expressing ligands for multiple T cell repressor receptors. While the functions of T cell inhibitory receptors such as CTLA4, PD1 and BTLA have been well studied and the impact of engaging these receptors for modulating T cell function has been recognized^1,14,17,23^, our study describes a DC-based approach to hyper-activate these T cell inhibitory pathways individually or in combination during antigen presentation to induce robust T cell tolerance.

We predicted that engineering the DCs to ectopically / over express T cell inhibitory ligands will result in selective and enhanced engagement of negative regulatory-receptors on T cells, especially during antigen specific activation, and modulate the function of these T cells resulting in antigen specific immune tolerance. Since early activation of T cells results in increased expression of negative regulatory receptors such as CTLA4, PD-1 and BTLA^13,17,19,23,40^, enhanced engagement and active T cell repression can occur mainly in antigen experienced / TCR engaged T cells. Further, since the natural expression of T cell activation ligands such as CD80 and CD86 on these engineered DCs is not restricted due to ectopic expression of T cell inhibitory receptor selective ligands, they can activate T cells initially to provide antigen specificity and to upregulate repressor-receptor expression. It will then be followed by a strong suppression of these T cells mediated by enhanced-ligation induced active signaling through these receptors. Validating this notion, here we show that LPS-treated (mature) DCs that are ectopically expressing CTLA4, PD1 and BTLA selective ligands, primarily aimed at enhanced inhibitory receptor engagement upon antigen presentation, can effectively induce CD4+ T cell hyporesponsiveness both in vitro and in vivo. Moreover, we show that treatment with DCs that ectopically express selective ligands of all three receptors results not only in the modulation of both T and B cell responses in vivo, but also ameliorates disease severity in a self-antigen immunization induced autoimmune model.

Mature DCs such as those treated with LPS are potent APCs that express high levels of antigen presentation associated surface molecules such as MHC, CD80 and CD86 as well as various cytokines^44,48^; hence ideal for activating T cells in an antigen specific manner. Further, mature DCs, as compared to immature DCs, could be less susceptible to microenvironment induced pro-inflammatory changes and not expected to produce undesired outcomes. In addition, unlike endogenous proteins, microenvironment is not expected to profoundly influence the levels of ectopically expressed surface ligands. Therefore, CTLA4, PD1 and BTLA selective ligand expressing mature DCs can produce more reliable and consistent outcomes when used in vivo for inducing antigen specific immune modulation. Our results showing similar trends in the outcomes, when untagged and fluorescent protein tagged T cell negative regulatory ligand expressing DCs, with two antigens and in two strains of mice were used, appear to validate this notion. Of note, lentiviral transduction and LPS activation associated expression of secreted factors and cell surface molecules by ligand overexpressing DCs could contribute to modulation of T cell properties differently than that by APC subsets that naturally express elevated levels of these ligands. This is evident from our observation, in contrary to previous reports^21,49–54^, PD-L1 overexpressing engineered DCs do not induce an increase in Foxp3+ T cells, but these DCs suppress both Th1 and Th17 (IFNγ and IL17) responses. Nevertheless, the overall impact of engineered mono- and multi-ligand DCs on T cell function, as anticipated with enhanced engagement of T cell repressor receptors, is hyporesponsiveness/tolerance.

Our in vitro and in vivo studies using mono-ligand and multi-ligand DCs, as anticipated, show that while enhanced CTLA4, PD1 and BTLA engagements by engineered DCs during antigen presentation result in an overall suppression of T cell proliferation and IL2 production, key differences in the type and magnitude of immune responses are evident upon targeting specific receptors. While enhanced CTLA4 engagement by B7.1wa DCs induced TGFβ1 and IL10, enhanced PD1 engagement induced only IL10, but not TGFβ1. Further, increase in Foxp3+ T cells appeared to be associated only with enhanced CTLA4 engagement, but not PD1 or BTLA engagement by respective DCs. In fact, earlier reports including ours ^32,34–36^ have shown that enhanced CTLA4 engagement up on antigen presentation can induce similar IL10, TGFβ1 and Foxp3 producing/expressing cell responses. However, in contrary to previous reports on the impact of PD-1 signaling in Treg induction^49–53^, engineered PD-L1 DCs did not show an increase in Foxp3+ cells in vitro or in vivo, perhaps due to higher expression of pro-inflammatory cytokines and activation markers by these virus-transduced LPS-treated DCs. Albeit the differences in the magnitude and type of responses, immune modulation by B71.wa, PDL1 and HVEM-CRD1 DCs involved diminished IFNγ response by T cells and an overall suppression of antibody production. Importantly, enhanced CTLA4 engagement by B7.1wa DCs caused enhanced IL17 response, perhaps influenced by higher TGFβ1 production by CTLA4 engaged CD4+ T cells. However, such an IL17 response was profoundly suppressed by multi-ligand DCs, which includes B7.1wa, suggesting a more efficient suppression of pro-inflammatory response upon enhanced engagement of multiple T cell repressor receptors. Superior suppression of T cell proliferation, IL2, IFNγ, as well as antigen-specific antibody responses, and the maintenance of Foxp3+ T cell and IL10 response inducing abilities of individual ligand DCs by multi-ligand DCs substantiates this notion. Of note, previous reports have shown that DCs that express higher levels of PD-L1 suppress Th1 response, but not Th17 response. This discrepancy is perhaps associated with the type, maturation status, and PD-L1 levels of the DCs. Importantly, although B7.1wa DCs induce higher IL17 response, profoundly suppressed autoantibody and thyroiditis severity in EAT mice treated with these DCs suggest that this, especially in association with robust Treg and IL10 responses, is not a pathogenic response. Furthermore, many previous studies including ours^55–59^ have shown that Th17/IL17 response in association with increased Tregs promotes protection from the disease in other autoimmune models.

Although CTLA4, PD1 and BTLA have inhibitory/repressor functions in T cells as indicated by suppression of IL2 production and T cell proliferation^13,17,19,23,40^, signaling by these receptors can produce different immune outcomes. We and others have shown that CTLA4 engagement during antigen presentation can induce both TGFβ1 and IL10 expression in T cells and promote Foxp3+ and IL10+ Treg generation^32,34–36,60^. PD1 activation has been shown to result in increased IL10 and Foxp3 expressing T cells^61–64^. BTLA activation also contributes to generation of IL10 expressing T cells^65,66^. Reciprocally, activation of these signaling pathways often suppresses the production of pro-inflammatory cytokines. Importantly, as indicated by the diminished levels of antigen specific antibodies in treated mice, our observations show that T cell inhibitory ligand expressing engineered DC induced modulation of T cell responses can have profound impact on B cell function. While overexpression of selective ligands for one or multiple T cell repressor receptors on mature DCs that already express elevated levels of costimulatory molecules and a multitude of cytokines can add to the complexity of DC-T cell interactions and the immune outcomes, our observations concur with the previous known features of CTLA4, PD1 and BTLA engagement associated outcomes to a large extent. The differences in immune response when different ligand DCs were used can be in part also due to activation of different downstream signaling pathways by these receptors in T cells. However, previously reported phenomena such as 1) reverse signaling through the ligands expressed on APCs, 2) competition with other receptors or binding to more than one ligand, and 3) unique mechanism such as trans-endocytosis of ligands^3,12,67^ may also influence the type and magnitude of overall T cell response induced by different engineered DCs.

Treg induction and immune tolerance can occur under various conditions including microbial infections, and upon cytokine and antibody therapies^32,34,35,49,53,56,68,69^. DCs often play a central role in inducing Tregs under these conditions, substantiating the notion that DCs are crucial for T cell activation as well as regulation. These contrasting properties are dependent on maturation status and/or the cytokine and surface molecular expression profiles of these cells. Higher PD-L1 expression and IL10, and lower costimulatory molecule expression have been linked to the ability of DCs to induce Tregs^49,50,53,68,70–74^. Treatment using factors such as cytokines and normal immunoglobulin can promote tolerogenic properties / immature phenotype of DCs resulting in Treg induction in vivo^68,73,75,76^. Importantly, while mature DCs are usually linked to immune activation rather than tolerance, here we show that robust Treg and/or T cell tolerance can be achieved using T cell inhibitory ligand expressing engineered mature DCs. While the tolerogenic properties of these DCs are dependent on the expression levels of T cell inhibitory receptor selective ligands, mature phenotype provides these cells with the ability to present antigens effectively and induce antigen specific Tregs and/or tolerance.

Overall, our observations show that powerful tAPCs can be generated by engineering the DCs to express selective ligands of T cell repressor receptors, and this could be an attractive and effective approach for antigen specific autoimmune therapy. Moreover, the ability of these engineered DCs, multi-ligand DCs particularly, to affect multiple aspects of immune function, viz: higher Treg frequencies and immune regulatory cytokine responses, and lower pro-inflammatory IFNγ responses as well as both IgG2a and IgG1 responses against autoantigen suggests that such tAPCs could have therapeutic value in both T cell mediated and antibody mediated autoimmune diseases. Challenges of employing DC based approaches for human autoimmune therapy include engineering the DCs ex vivo efficiently and consistently for therapy and maintaining the tolerogenic phenotype and function of DCs in vivo upon therapeutic injection. As stated above, efficient engineering of the DCs for constitutive overexpression of multiple T cell inhibitory ligands, as described in this study, makes them less susceptible to changes in the functional properties and produces undesired outcomes upon in vivo therapeutic delivery; hence highly desired for clinical translation for treating autoimmune diseases such as thyroiditis and type 1 diabetes where self-antigens are known. Induction of antigen specific T cell tolerance is the key feature of DC based tolerogenic approaches. Although this approach needs to be further evaluated in a spontaneous model of autoimmunity, based on the outcomes of this proof-of-principle study, we envision that DCs generated from the human peripheral blood monocytes can be engineered similarly to express human T cell inhibitory ligands, loaded with antigen(s) of interest, and these autologous DCs will be injected back to the patient to reliably suppress the autoimmunity. While suppression of autoimmunity in at-risk subjects at pre-clinical stages can effectively prevent the clinical onset of disease, treatment of subjects with established diseases could result in minimized autoimmunity associated disease complications. Importantly, although mono-ligand DCs alone could be effective in inducing T cell tolerance in humans, our pre-clinical observations suggest that engineered multi-ligand DC therapy will be more robust in suppressing autoimmunity and associated complications.

## Materials and methods

### Mice, antigens, antibodies and other reagents

C57BL6, CBA/J, IL10−/− and OT-II TCR-transgenic (TCR-Tg) mice (OT-II mice) were originally purchased from the Jackson laboratory (Bar Harbor, ME, USA). Foxp3-GFP-knockin (ki)^77^ mice in the B6 background were kindly provided by Dr. Kuchroo (Harvard Medical School, MA). To generate OT-II-TCR-Tg-Foxp3-GFP-ki (OT-II-Foxp3-GFP) mice, OT-II mice were crossed with Foxp3-GFP-ki mice. Breeding colonies of these mice were established and maintained in the specific pathogen free facilities of the University of Illinois at Chicago (UIC) and Medical University of South Carolina (MUSC). All experimental protocols were approved by the Institutional Animal Care and Use Committees (IACUC) of UIC and MUSC. All methods in live animals were carried out in accordance with relevant guidelines and regulations of these committees. Details of other reagents and sources are listed in Supplemental Table 1.

### Cloning and lentivirus production

The third-generation replication-incompetent lentiviral cDNA cloning vector and packaging plasmids (Supplemental Fig. 1) were purchased from SBI and modified for this study. For cloning eCFP, eYFP and mRFP cDNA, pCDH1-copGFP vector was digested using NheI (downstream of CMV promoter) and SalI (downstream of copGFP) restriction enzymes, and the restriction digestion fragment was replaced with PCR amplified (from various vectors) and NheI and SalI digested eCFP, eYFP and mRFP cDNAs as depicted in Fig.1. The final plasmids pCDH1-eCFP, pCDH1-eYFP, and pCDH1-mRFP, and the original plasmid pCDH1-copGFP were used as control vectors. WT PD-L1 cDNA was amplified from mouse spleen cells. For generating B7.1wa cDNA, ectodomain region was amplified from VR1255-B7.1wa-Ig vector^78^ (kindly provided by Dr. Gérald J. Prud’homme, Toronto, Canada), and fused with transmembrane/cytoplasmic domains of B7.1 by overlap PCR. HVEM amino-terminal CRD1^24^ and the transmembrane/cytoplamic tail regions were PCR amplified separately and fused by overlap PCR to generate HVEM-CRD1 cDNA. These cDNA were cloned into aforementioned original pCDH1-copGFP plasmid between NheI and NotI sites, or into pCDH1-eCFP, pCDH1-eYFP or pCDH1-mRFP plasmids between NheI and XhoI as fusion proteins at the N-terminus of fluorescent tag. cDNAs of all vector constructs were sequence confirmed before use. For lentivirusproduction, HEK293T cells were transfected with above cDNA expression vectors along with packaging vectors by calcium phosphate method. Occasionally, GPRG cells^79^, provided by National Gene Vector Biorepository (Indianapolis, IN, USA), were used as virus packaging cells. Virus-containing media were collected after 36 and 72 h, pooled and centrifuged at 3500 RPM for 15 min and the supernatants were subjected to 0.22-μm filtration, and concentrated by ultracentrifugation, high speed centrifugation for 48 h, or PEG precipitation. Infectious titers/transduction unit (TU) of the virus were determined by using fresh HEK293T cells before using for transduction of DCs.

### Generation of BM derived DCs

DCs were generated *in vitro* from BM cells as described before^34–36^. Briefly, BM cells were cultured in complete RPMI 1640 medium containing 10% heat-inactivated FBS in the presence of GM-CSF (20 ng/ml) at 37°C in 5% CO_2_ for 3 days and then for further 3 days in fresh complete RPMI 1640 medium containing GM-CSF (20 ng/ml) and IL-4 (5 ng/ml). For lentiviral transduction, cells from 3 day old cultures were used. A final concentration of > 5×10^8^ TU/ml virus was used for transducing DCs in the presence of polybrene (8 μg/ml) and protamine sulfate (10 μg/ml) for up to 24 h, washed and cultured in fresh medium with GM-CSF and IL4 for an additional 48 h. For some experiments, bacterial LPS (Sigma) was added to these cultures (1.0 μg/ml) at 48 h post-viral transduction and cultured for an additional 24 h. Transduction efficiency was examined by microscopy and FACS, and the cells that showed transduction efficiency of >90% were washed thoroughly and counted before in vitro or in vivo use.

### Phagocytic assay

Lentivirus (CFP control vector) transduced and non-transduced DCs were left untreated or exposed to LPS for 24 h, incubated with varying number of 1 μm yellow fluorescent latex beads (Polysciences Inc) for 2 h or 6 h, stained with CD11c specific Ab and analyzed by FACS.

### In vitro antigen presentation assay

Ova-MHC class II peptide pulsed DCs (2 × 10^4^ cells/well) were plated in triplicate in 96-well U-bottom tissue culture plates along with unlabeled or CFSE-labeled purified CD4+ cells from OT-II or OT-II-Foxp3-GFP mice (1 × 10^5^ cells/well) in complete RPMI 1640 medium. Control antibody, anti-CTLA4 antibody, PD1-Ig, BTLA-Ig or anti-TGF-β1 antibody were added to some cultures. After 4 days of culture, cells were subjected to surface and/intracellular staining using fluorochrome labeled Abs and examined for CFSE dilution or cytokine production by FACS. Cells were subjected to brief activation using PMA and ionomycin (4 hours) in the presence of Brefeldin A before staining to detect intracellular cytokines. In some assays, equal number of cells from primary cultures were seeded along with OT-II peptide, freshly isolated splenic DCs for 24 h and the spent media was tested for cytokine levels by ELISA or multiplex assays.

### Ova and mTg priming, and treatment with engineered DCs

For inducing Ova specific immune response, six to eight week old C57/BL6 mice were injected (i.v.) with Ova along with bacterial LPS (10 μg Ova and 5 μg LPS) on days 0 and 10. For inducing EAT, 8 week old CBA/J mice were i.v. injected with 100 μg of mTg along with 25 μg of bacterial LPS on days 0 and 10. Engineered DCs were cultured (2×10^6^ DCs/ml) for last 24 h in the presence of antigen (10 μg Ova or 25 μg mTg/ml) and bacterial LPS (1 μg/ml), washed, and injected i.v. (2 × 10^6^ DCs/mouse) twice at a 10-day interval (on days 5 and 15). Ova primed mice were euthanized between 25 days post-priming to determine immune response. Ova specific T cell proliferative and cytokine responses were examined using spleen by ELISA or multiplex assay, and serum samples were tested for Ova specific Ab response by ELISA as described earlier^33,35,80^. Mice that received mTg for inducing EAT were euthanized between 30 days of priming for determining T cell response using spleen and cervical LN cells, or on day 40 for examining thyroiditis severity and serum autoantibody levels.

### Detection of anti-Ova and mTg Abs

Serum levels of anti-Ova or anti-mTg-specific total IgG, IgG1 and/or IgG2a Abs were determined by ELISA. Ninety six-well plates were coated with 0.2 μg/well Ova or 0.5 μg/well mTg (100 μl) in 0.01 M carbonate-bicarbonate buffer, pH 9.6, overnight at 4°C. Wells were blocked with 300 μl of PBS containing 1% BSA for 1 h at RT. Serial dilutions of serum samples were added to the wells in triplicate and incubated for 1 h at RT. After washing, the plates were further incubated with HRP-labeled anti-mouse IgG, IgG1, or IgG2a for 1 h at RT. The enzyme reaction was developed using TMB/hydrogen peroxide substrate and the absorbance was measured at 450 nm. Highest dilution that showed OD value ≥0.05 above the background OD was considered the antibody titer of a given sample.

### Evaluation of EAT

Thyroids along with trachea were removed and fixed in 4% formaldehyde, 5-μm paraffin sections were made, and subjected to H&E staining to examine lymphocyte infiltration. Cellular infiltration and thyroid follicular damage were scored as grades 0 −4 as described in our earlier reports33,35,80.

### Statistical analysis

Mean, SD, and statistical significance (*p*-value) by paired or unpaired T cells were calculated using a Microsoft Excel or GraphPad prism statistical application. In most cases, values of the individual test group (ligand-DC group) were compared with that of control group (control-DC group). Statistical analysis of thyroiditis severity was done by Fisher’s exact test. *p* ≤ 0.05 was considered significant.

## Author contribution

R.G. researched and analyzed data and edited the manuscript, S.K. researched data, N.P. researched data, G.L. researched data and C.V. designed experiments, researched and analyzed data, and wrote/edited manuscript.

## Data availability

The datasets generated during and/or analyzed during the current study are available from the corresponding author on reasonable request.

## Resource availability

The cDNA vectors generated during the current study is available from the corresponding author on reasonable request

## Footnote

This work was supported by unrestricted research funds from MUSC and National Institutes of Health (NIH) grants R21AI069848 and R01AI073858 to C.V. C.V. is the guarantor of this work and, as such, has full access to all the data in the study and takes responsibility for the integrity of the data and accuracy of the data analysis. The authors are thankful to Cell and Molecular Imaging, Pathology, Proteomics, immune monitoring and discovery, and flow cytometry cores of MUSC and UIC for the histology service, microscopy, FACS and multiplex assay instrumentation support.

