## Supplemental data for "Engineered Dendritic Cell-Directed Concurrent Activation of Multiple T cell Inhibitory Pathways Induces Robust Immune Tolerance"

and Chenthamarakshan Vasu

Supplemental Table 1:

| Item | Vendor | Catalog # |
|---|---|---|
| Chicken ovalbumin | Sigma-Aldrich, St. Louis, MO, USA | A7641 |
| pCDH-CMV-MCS-EF1α-copGFP | System Biosciences, Mountain View, CA,USA | CD511A-1 |
| Lentivector Packaging Kit | System Biosciences, Mountain View, CA, USA | LV500A-1 |
| pcDNA3-eCFP | Addgene, Watertown, MA, USA | 13030 |
| pcDNA3-eYFP | Addgene, Watertown, MA, USA | 13033 |
| pcDNA3-mRFP | Addgene, Watertown, MA, USA | 13032 |
| Anti-mouse CD16/CD32 | BD Biosciences, San Jose, CA, USA | 553140 |
| Anti-mouse CD11c-PEcy5 | BD Biosciences, San Jose, CA, USA | 553802 |
| Anti-mouse CD4-PEcy5 | BD Biosciences, San Jose, CA, USA | 553050 |
| Anti-mouse CD4 –PE-PETR | Invitrogen, Carlsbad, CA, USA | MCD0417 |
| Anti-mouse CD25-Alexa488 | Invitrogen, Carlsbad, CA, USA | RM6004 |
| Anti-mouse IFN-γ-PE | eBiosciencs/Affymetrix, Santa Clara, CA | 12-7311-82 |
| Anti-mouse IFN-γ-Alexa488 | eBiosciencs/Affymetrix, Santa Clara, CA | 53-7311-82 |
| Anti-mouse IL-17A-Alexa488 | eBiosciencs/Affymetrix, Santa Clara, CA | 53-7177-81 |
| Anti-mouse IL-17A-PE | eBiosciencs/Affymetrix, Santa Clara, CA | 12-7177-81 |
| Anti-mouse TNF-α-PE | eBiosciencs/Affymetrix, Santa Clara, CA | 12-7321-82 |
| Anti-Mouse IL-10-PE | eBiosciencs/Affymetrix, Santa Clara, CA | 12-7101-82 |
| Anti-Mouse IL-10-Alexa488 | eBiosciencs/Affymetrix, Santa Clara, CA | 53-7101-82 |
| Anti-mouse CD80-PE | BD Biosciences, San Jose, CA, USA | 562504 |
| Anti-mouse CD80-FITC | BD Biosciences, San Jose, CA, USA | 561954 |
| Anti-mouse CD86-PE | BD Biosciences, San Jose, CA, USA | 553692 |
| Anti-mouse CD40-PE | BD Biosciences, San Jose, CA, USA | 553658 |
| Anti-mouse I-$A^d$ -PE | BD Biosciences, San Jose, CA, USA | 553548 |
| Anti-mouse I-$A^b$ -PE | BD Biosciences, San Jose, CA, USA | 562012 |
| Anti-mouse I-$A^b$ -PE | BD Biosciences, San Jose, CA, USA | 553537 |
| Anti-mouse PD-L1-PE | eBiosciencs/Affymetrix, Santa Clara, CA | 12-5982-82 |
| Anti-mouse HVEM-PE | BD Biosciences, San Jose, CA, USA | 12-5962-80 |
| Anti-mouse HVEM-Alexa488 | RnD Systems, Minneapolis, MN 55413 | FAB2516G |
| Anti-mouse CTLA4-PE | BD Biosciences, San Jose, CA, USA | 561718 |
| Anti-mouse CTLA4 | BD Biosciences, San Jose, CA, USA | 553718 |
| Anti-mouse PD1 | Biolegend, San Diego, CA 92121 | 135202 |
| Anti-mouse BTLA | Biolegend, San Diego, CA 92121 | 139104 |
| Anti-mouse Foxp3-PE | eBiosciencs/Affymetrix, Santa Clara, CA | 12-5773-82 |
| Anti-mouse Foxp3-PE-Cy5 | eBiosciencs/Affymetrix, Santa Clara, CA | 15-5773-82 |
| Anti-mouse IL-10 | eBiosciencs/Affymetrix, Santa Clara, CA | 16-7101-81 |
| CTLA4-Ig | RnD Systems, Minneapolis, MN 55413 | 434-CT-200/CF |
| PD1-Ig | RnD Systems, Minneapolis, MN 55413 | 1021-PD-100 |
| BTLA-Ig | RnD Systems, Minneapolis, MN 55413 | 3007-BT-050 |
| Anti-mouse-TGFβ1 | RnD Systems, Minneapolis, MN 55413 | MAB1835-500 |
| Anti-mouse LAP-APC | Biolegend, San Diego, CA 92121 | 141405 |
| CD4+ T cell enrichment kit | Invitrogen, Carlsbad, CA, USA | 11416D |

| | | |
|---|---|---|
| CD4+ T cell enrichment kit | Miltenyi Biotec, Bergisch Gladbach, Germany | 130-104-454 |
| Mouse IL10 ELISA | eBiosciencs/Affymetrix, Santa Clara, CA | 88-7105-86 |
| Mouse TGFβ1 ELISA | RnD Systems, Minneapolis, MN 55413 | DY1679 |
| Mouse IL17A ELISA | eBiosciencs/Affymetrix, Santa Clara, CA | 88-7371-22 |
| Mouse IFN-γ ELISA | eBiosciencs/Affymetrix, Santa Clara, CA | 88-7314-22 |
| Mouse IL-12 ELISA | eBiosciencs/Affymetrix, Santa Clara, CA | 88-7121-22 |
| Mouse IL-1β ELISA | eBiosciencs/Affymetrix, Santa Clara, CA | 88-5019-22 |
| Mouse IL-2 ELISA | RnD Systems, Minneapolis, MN 55413 | DY402 |
| Mouse cytokine multiplex Assay | Invitrogen, Carlsbad, CA, USA | EPXR360-26092-901 |
| Mouse GM-CSF | Invitrogen, Carlsbad, CA, USA | PMC2011 |
| Mouse IL-4 | Invitrogen, Carlsbad, CA, USA | PMC0046 |
| PEG6000 | Sigma-Aldrich, St. Louis, MO, USA | 8074915000 |
| Polybrene | Sigma-Aldrich, St. Louis, MO, USA | S2667 |
| Protamine Sulfate | Sigma-Aldrich, St. Louis, MO, USA | P4020 |
| Bacterial LPS | Sigma-Aldrich, St. Louis, MO, USA | L7770 |
| Fluorescent latex beads | Polysciences, Warrington, PA, USA | 19121 |
| CFSE | Invitrogen, Carlsbad, CA, USA | V12883 |
| PMA | Sigma-Aldrich, St. Louis, MO, USA | P8139 |
| Ionomycin | Sigma-Aldrich, St. Louis, MO, USA | I0634 |
| Brefeldin A | BD Biosciences, San Jose, CA, USA | 555029 |
| 96 Well MaxiSorp plate (Nunc) | Thermo-Fisher, Waltham, MA | 44-2404-21 |
| Anti-mouse IgG-HRP | Invitrogen, Carlsbad, CA, USA | 61-6520 |
| Anti-mouse IgG1-HRP | Invitrogen, Carlsbad, CA, USA | A10551 |
| Anti-mouse IgG2a HRP | Invitrogen, Carlsbad, CA, USA | 61-0220 |
| Anti-human Fc-PE | Invitrogen, Carlsbad, CA, USA | H10104 |
| Anti-human Fc-Alexa488 | Invitrogen, Carlsbad, CA, USA | H10120 |
| TMB/peroxide substrate | Invitrogen, Carlsbad, CA, USA | 34028 |

### Supplemental fig. 1

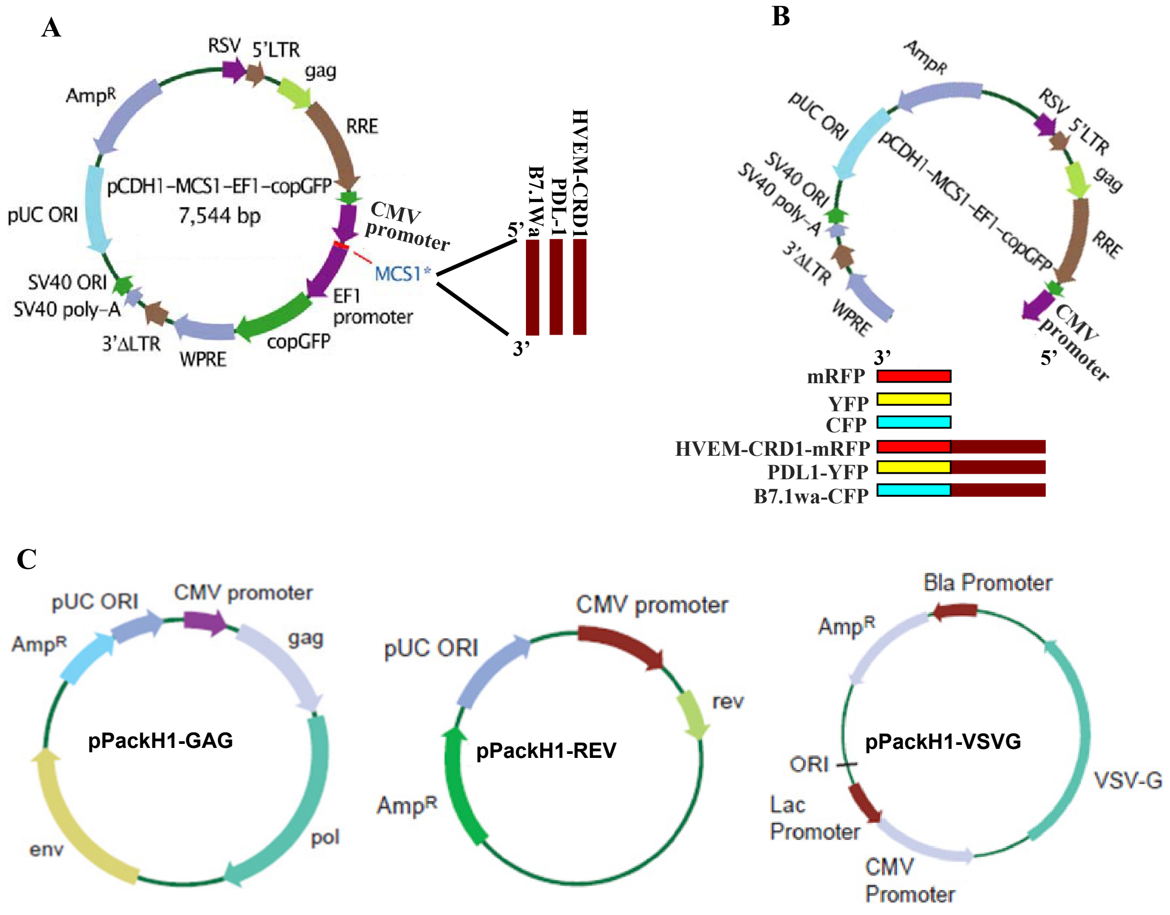


**Supplemental Fig. 1: Lentiviral system and the constructs used for this study. A)** T cell negative regulatory ligands were cloned individually without fluorescent tags under the CMV promoter of a third generation lentiviral vector from SBI Inc. In these vectors, GFP is expressed under a different (EF1) promoter. Unmodified vector was used as control. **B)** Ligands were also cloned as fluorescent protein (CFP, YFP and RFP) fusion. GFP and the associated promoter were deleted from these constructs. Vector constructs that express CFP, YFP and RFP separately were used as controls. **C)** Shown packaging vectors originally purchased as a pool from SBI were separated, propagated and used for generating lentivirus in 293T cells or GPRG cells. Original or edited vector maps from SBI Inc are shown.

### Supplemental fig. 2

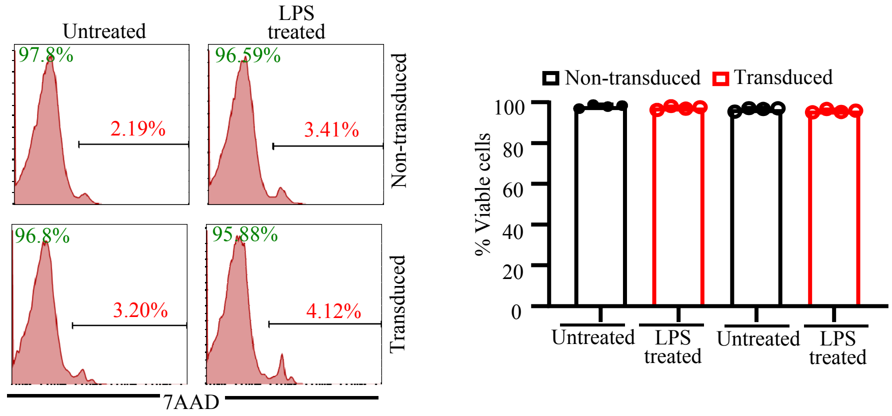


**Supplemental Fig. 2: Viability of lentivirus transduced and LPS treated DCs.** C57/BL6 BM DCs were transduced with lentiviral vectors and incubated with bacterial LPS for 24 h as described in Fig. 1. Cells were harvested and examined for viability by flow cytometry after staining with 7-AAD. Representative FACS graphs (left panel) and mean±SD of 7-AAD negative (live) cell frequencies from 3 parallel transductions (right panel) are shown.

### Supplemental fig. 3

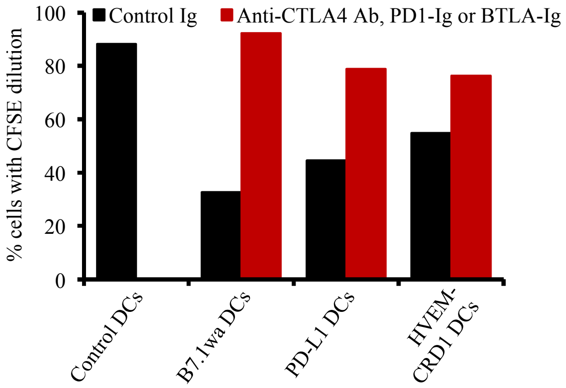


**Supplemental Fig. 3: Ligand specificity of engineered DC mediated suppression of T cell proliferation upon antigen presentation.** Antigen presentation/CFSE dilution assay was conducted as described in, and in parallel with the experiment of, Fig. 2C. Culture wells specific to this figure were added either with control Ig, or respective receptor blocking antibody (anti-CTLA4 Ab) or soluble receptor (PD1-Ig or BTLA-Ig). Mean values of 2 independent experiments, done in triplicate, using parallel preparations of DCs are shown. This experiment was repeated at least once with similar trend.

Supplemental fig. 4

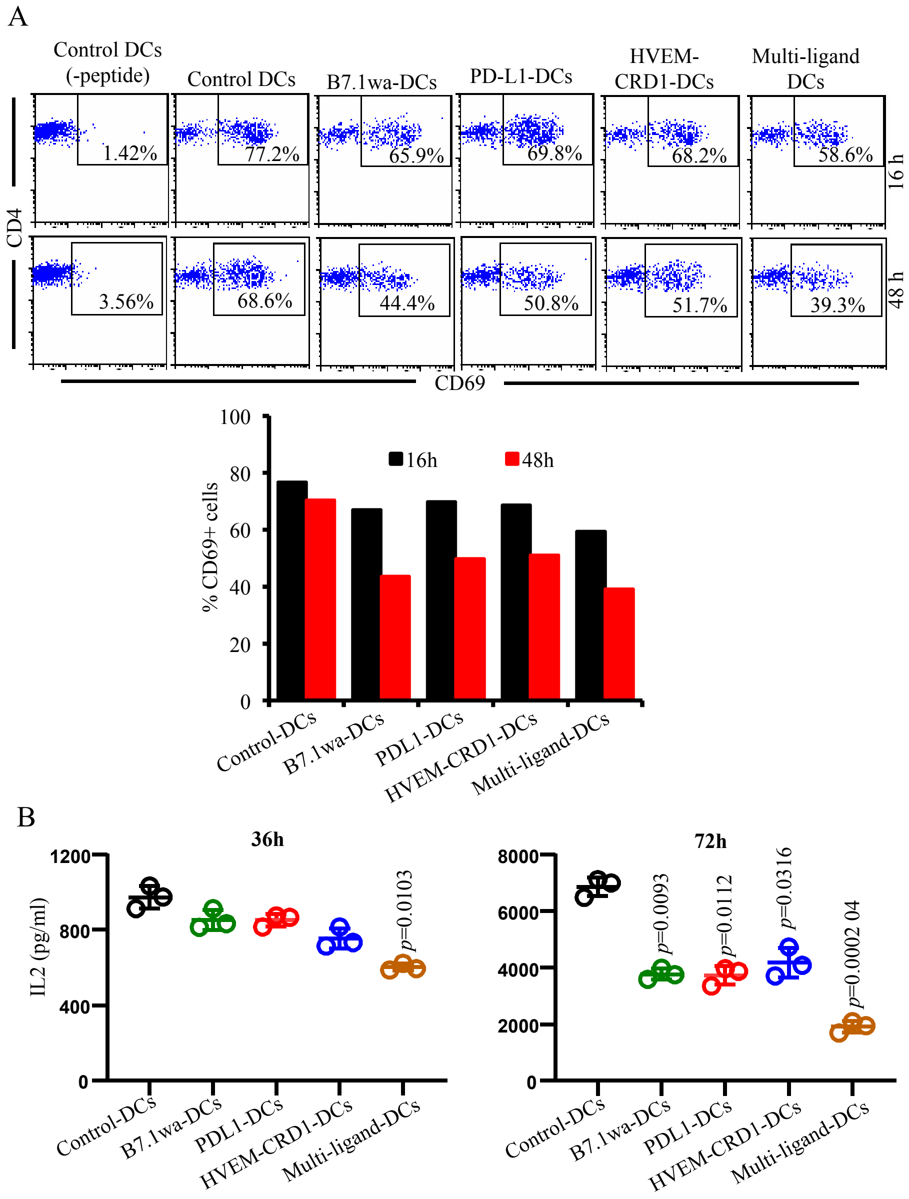


**Supplemental Fig. 4: T cell activation dynamics upon activation using T cell inhibitory ligand expressing engineered DCs.** Antigen presentation assay was conducted as described in Figs. 3C and 4A. **A)** Cells from primary cultures were harvested at 16 and 48 h time-points and examined for the frequencies of cells with CD69 expression by FACS. Representative FACS plots (upper panel), and mean values of 2 independent experiments, done in triplicate, using parallel preparations of DCs (lower panel) are shown. **B)** Supernatants from similar cultures were harvested at 36 and 72 h time-points and tested for IL2 levels by ELISA. Mean±SD values of 3 independent experiments, done in triplicate, using parallel preparations of DCs are shown. These experiments were repeated at least once with similar trend.

Supplemental fig. 5

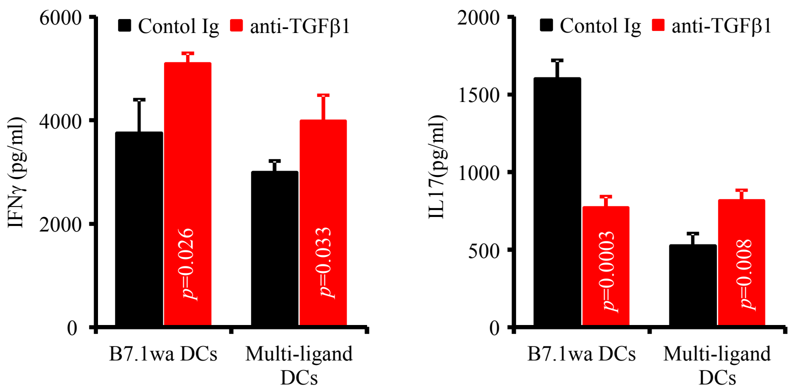


**Supplemental Fig. 5: IL17 response induced by B7.1wa DCs is due to TGFβ1 produced in the culture.** Supernatants of cultures described under Fig. 4C were examined for cytokine levels by ELISA. Mean±SD values of cytokine levels of 3 independent experiments, done in triplicate, using parallel preparations of DCs are shown. This assay was repeated once with similar trend. *p*-value by paired *t*-test. Cultures containing anti-TGFβ1 antibody were compared to respective control cultures.

Supplemental fig. 6

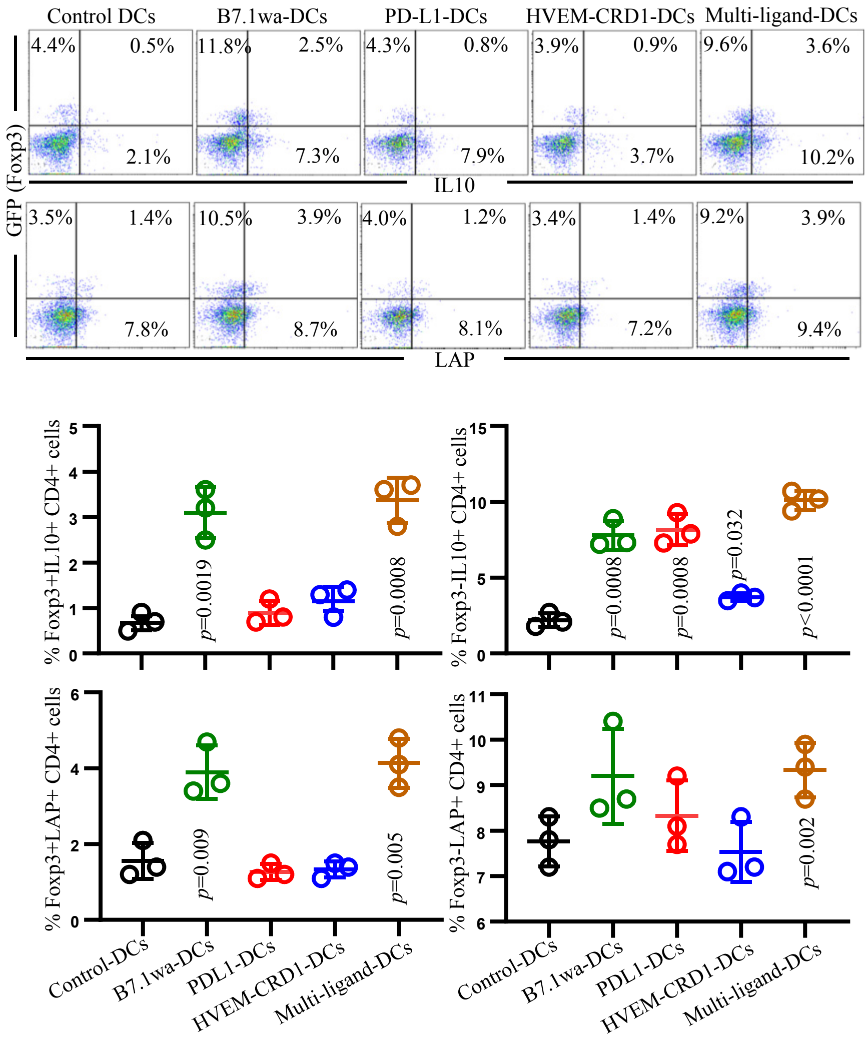


**Supplemental Fig. 6: IL10 and LAP expression by ligand DC activated CD4+ T cells.** Control and ligand DCs and OT-II-Foxp3-GFP T cells were used in antigen presentation assays as described for Fig. 4. Cells from primary cultures were stimulated using PMA and ionomycin in the presence of brefeldinA for 4 h, stained for surface CD4 and LAP and intracellular cytokines IL10, and subjected to FACS analysis. Representative FACS plots (upper panel) and mean±SD of Foxp3+IL10+, Foxp3-IL10+, Foxp3+LAP+ and Foxp3-LAP+ cell frequencies (among CD4+ T cells) of assays using 3 parallel preparations of DCs (lower panel) are shown. Each ligand DC group was compared separately with control DC group.

### Supplemental fig. 7

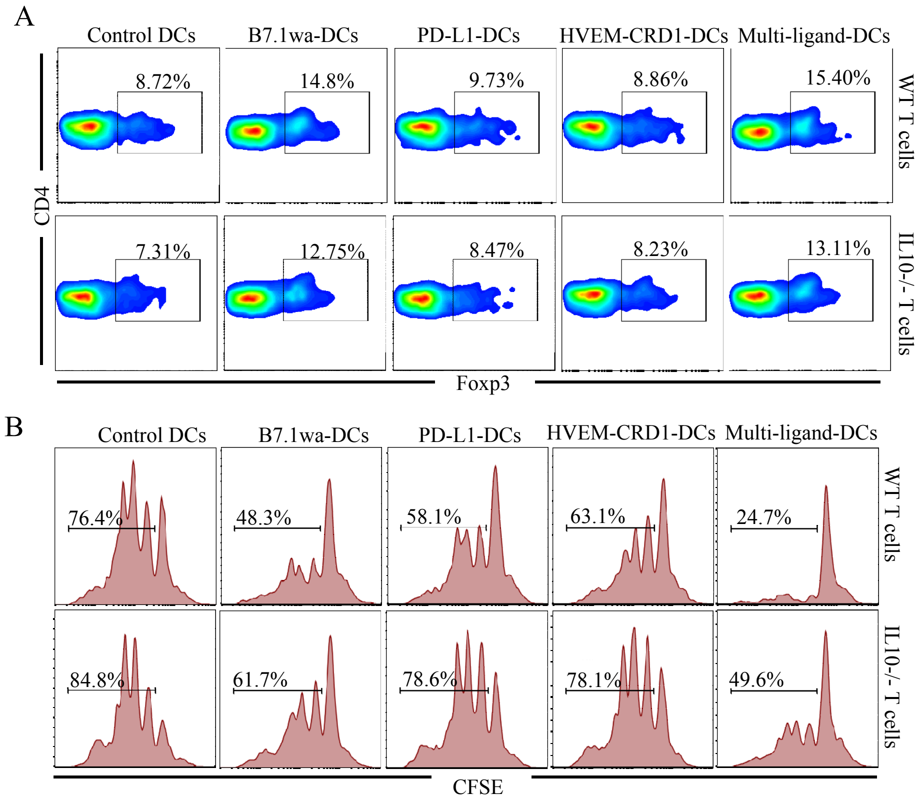


**Supplemental fig. 7: IL10 plays a key role in immune modulation by ligand DC activated CD4+ T cells.** CD4+ T cells isolated from WT and IL10-/- B6 mouse spleens were activated using anti-CD3 antibody in the presence of control and ligand DCs for 4 days. A) Cells were harvested and subjected to FACS analysis and Foxp3+ cells among CD4+ T cells were determined. B) In a co-culture assay, cells from these primary cultures were incubated in the presence of CFSE labeled fresh WT CD4+ T cells at 1:2 (effector : Treg) ratio, in plates that were coated with anti-CD3 and anti-CD28 antibodies, for 4 days and subjected to FACS. Percentage of CD4+ CFSE+ cells with CFSE dilution are shown. These results are representative of one of two independent experiments that produced comparable results. Each experiment was done in triplicate.

### Supplemental fig.8

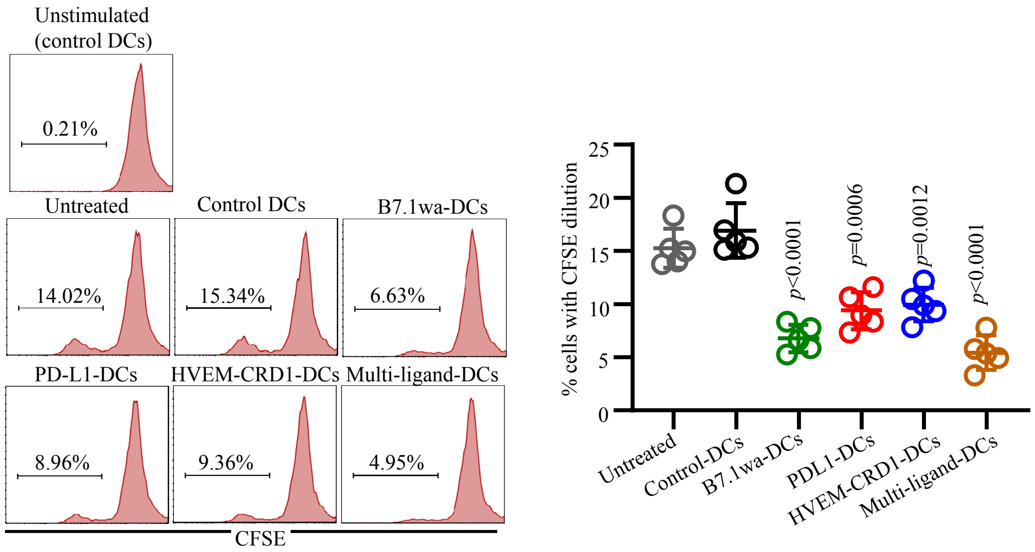


**Supplemental fig. 8: Treatment with T cell negative regulatory ligand expressing DCs diminishes mTg specific proliferative response of T cells.** Cervical LN cells from mice described in Fig. 6 were labeled with CFSE and cultured in the presence of mTg (5 mg/ml). Cells were harvested and subjected to FACS for determining the % of CD4+ T cells with CFSE dilution. Representative FACS plots (left panel) and Mean±SD (5 mice/group) of percentage of CD4+ T cells with CFSE dilution (right panel) are shown. Assays were performed using cells from individual mice in duplicate. *p*-value by unpaired *t*-test. Each ligand DC group was compared separately with control DC group.

Supplemental fig. 9

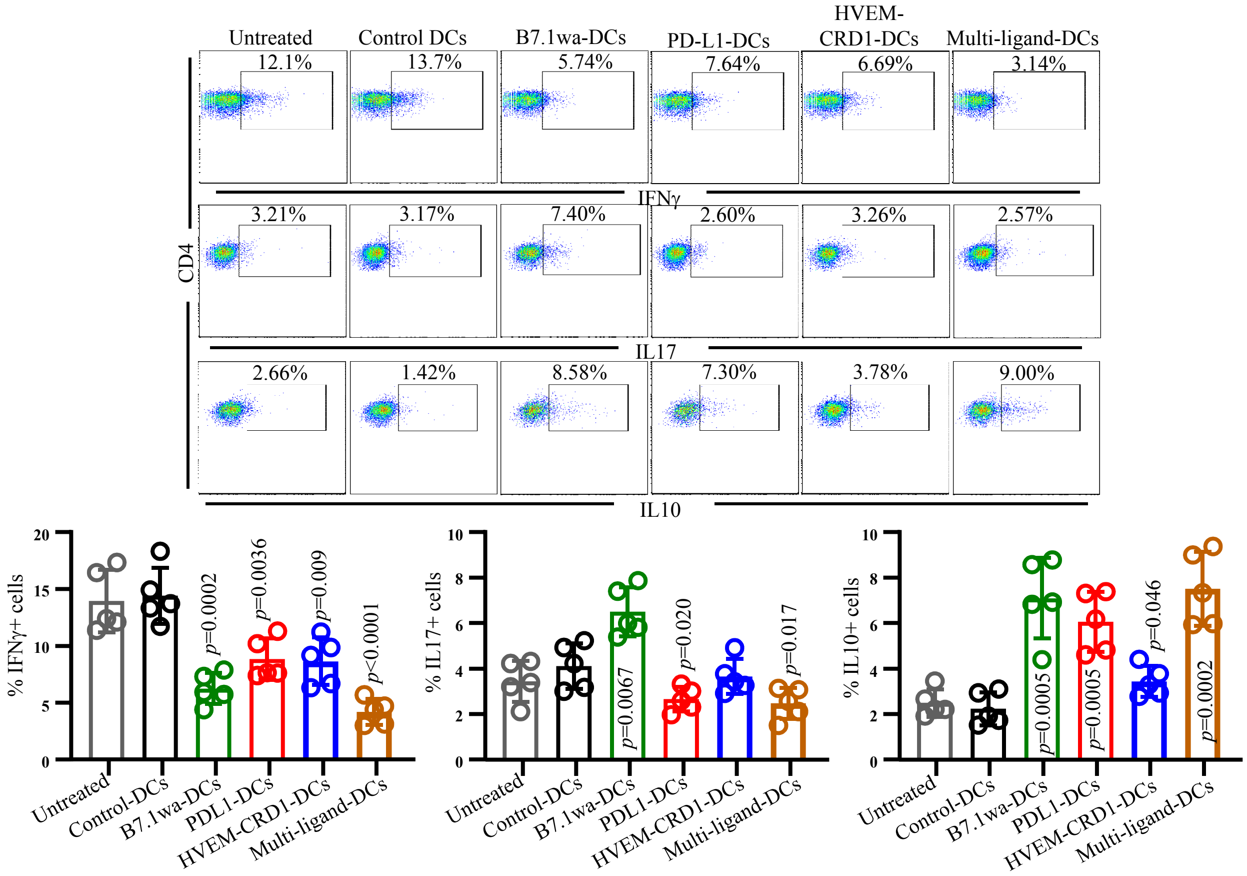


**Supplemental fig. 9: Treatment with T cell negative regulatory ligand expressing DCs results in modulation of T cell phenotype in EAT model.** Fresh spleen cells from mice described in Fig. 6 were subjected to brief (4 h) activation using PMA/Ionomycin in the presence of brefeldin A and subjected to intracellular staining for various cytokines. CD4+ population was gated for determining cytokine positive T cell frequencies. Representative FACS graphs (upper panel) and mean±SD of cytokine positive CD4 T cell frequencies of 5 mice/group (lower panel) are shown. *p*-value are by unpaired *t*-test. Each ligand DC group was compared separately with control DC group.
